# HLA-dependent variation in SARS-CoV-2 CD8+ T cell cross-reactivity with human coronaviruses

**DOI:** 10.1101/2021.07.17.452778

**Authors:** Paul R. Buckley, Chloe H. Lee, Mariana Pereira Pinho, Rosana Ottakandathil Babu, Jeongmin Woo, Agne Antanaviciute, Alison Simmons, Graham Ogg, Hashem Koohy

## Abstract

Pre-existing T cell immunity to SARS-CoV-2 in individuals without prior exposure to SARS-CoV-2 has been reported in several studies. While emerging evidence hints toward prior exposure to common-cold human coronaviruses (HCoV), the extent of- and conditions for-cross-protective immunity between SARS-CoV-2 and HCoVs remain open. Here, by leveraging a comprehensive pool of publicly available functionally evaluated SARS-CoV-2 peptides, we report 126 immunogenic SARS-CoV-2 peptides with high sequence similarity to 285 MHC-presented target peptides from at least one of four HCoV, thus providing a map describing the landscape of SARS-CoV-2 shared and private immunogenic peptides with functionally validated T cell responses. Using this map, we show that while SARS-CoV-2 immunogenic peptides in general exhibit higher level of dissimilarity to both self-proteome and -microbiomes, there exist several SARS-CoV-2 immunogenic peptides with high similarity to various human protein coding genes, some of which have been reported to have elevated expression in severe COVID-19 patients. We then combine our map with a SARS-CoV-2-specific TCR repertoire data from COVID-19 patients and healthy controls and show that whereas the public repertoire for the majority of convalescent patients are dominated by TCRs cognate to private SARS-CoV-2 peptides, for a subset of patients, more than 50% of their public repertoires that show reactivity to SARS-CoV-2, consist of TCRs cognate to shared SARS-CoV-2-HCoV peptides. Further analyses suggest that the skewed distribution of TCRs cognate to shared and private peptides in COVID-19 patients is likely to be HLA-dependent. Finally, by utilising the global prevalence of HLA alleles, we provide 10 peptides with known cognate TCRs that are conserved across SARS-CoV-2 and multiple human coronaviruses and are predicted to be recognised by a high proportion of the global population. Overall, our work indicates the potential for HCoV-SARS-CoV-2 reactive CD8^+^ T cells, which is likely dependent on differences in HLA-coding genes among individuals. These findings may have important implications for COVID-19 heterogeneity and vaccine-induced immune responses as well as robustness of immunity to SARS-CoV-2 and its variants.

## Introduction

After more than a year, the severe acute respiratory syndrome coronavirus 2 (SARS-CoV-2) pandemic remains a global health challenge and causes huge economic burden. SARS-CoV-2 virus gives rise to COVID-19 disease, which is characterised by a heterogenous clinical outcome ranging from asymptomatic infection to severe acute respiratory distress and death. The virus has proven to be dynamic, and the emergence of ‘variants of concern’ (e.g., the delta variant) challenges the existing mitigation strategies including vaccine rollouts ^1^.

Although disease morbidity is associated with several factors including age, sex and aberrant immune response; the mechanisms and factors underpinning the heterogeneity of disease are incompletely understood ^2^. Furthermore, reports of differential immune responses following vaccination have started to emerge, demonstrating prior SARS-CoV-2 infection can enhance COVID-19 vaccine response compared with naïve individuals ^3,4^. Indeed, many questions regarding the magnitude and robustness of immune response in disease, variants of concern and/or COVID-19 vaccination in different individuals remain open despite the great recent efforts.

Several studies^5,6^ have illustrated that the correlates of immunity to SARS-CoV-2 are implicated by the presence of pre-existing immunological memory conferred from cross-reactivity to other viruses. On the one hand such cross-reactivity could modulate disease severity, vaccine response and/or protection against SARS-CoV-2 and its variants via presence of antigen-specific memory T cells ^7^. Conversely, cross-reactivity may provoke immunopathology through mechanisms such as antibody-dependent enhancement of infection, with the potential for virus-induced autoimmune disease in years to come ^8^.

Coronavirus strains that infect humans belong to either alpha or beta genera. The alphacoronaviruses contain HCoV-229E and -NL63 while the four lineages of betacoronaviruses include HCoV-OC43 and -HKU1, SARS-CoV and -CoV-2, MERS-CoV and other viruses only identified in bats. HCoV-OC43, -HKU1, -NL63 and −229E strains are known to cause mild to moderate ‘common cold’ symptoms whereas MERS-, SARS-CoV-1 and −2 can cause severe respiratory tract disease and death. Previous natural and experimental infection studies in humans suggest antibody cross-reactivity within-but minimal reactivity between-endemic human alpha and beta coronaviruses. Unlike antibodies, T cell cross-reactivity to SARS-CoV-2 appears to be more prevalent. Several recent studies have reported existence of SARS-CoV-2-specific T cells in unexposed individuals ^9–15^, although it appears that T cell cross reactivity is more pronounced in CD4^+^ than CD8^+^ T cells in these subjects.

Recent studies have provided varying insights regarding the presence of pre-existing CD8+ T cell immunity to SARS-CoV-2 conferred by HCoV. In an investigation into the immunodominant SARS-CoV-2 SPR* epitope – associated with HLA-B*07:02 – Nguyen et al^16^., found little evidence of cross-reactive exposure in pre-pandemic Australian samples. On the other-hand, Francis et al^15^., found evidence of pre-existing memory CD8+ T cells in naïve samples and have shown that HLA genotype conditions pre-existing CD8+ T cell memory to SARS-CoV-2, and they suggest that unexposed individuals with specific HLA alleles (such as HLA-B*07:02), may be more likely to possess cross-reactive memory T cells specific for the SPR* SARS-CoV-2 epitope. These disparate results may stem from differences in regional HLA allele frequencies and / or experimental methodology. Nevertheless, the extent to which patients’ haplotypes and SARS-CoV-2-HCoV cross-reactivity - amongst other factors - are linked to heterogeneous COVID-19 disease, robustness of immunity against SARS-CoV-2 and its variants, and/or protection after vaccine-induced immune response, remains to be elucidated.

In this study, we examined the evidence for SARS-CoV-2-specific T cell cross-reactivity with common-cold HCoVs and identifying 126 immunogenic SARS-CoV-2 peptides that are highly similar to 285 predicted HCoV pMHC. We additionally identified a set of SARS-CoV-2 peptides with high similarity to several human genes. We found that public TCR repertoires reactive to SARS-CoV-2 in COVID-19 patients who carry specific HLA alleles primarily recognise SARS-CoV-2 peptides with high similarity to HCoVs, suggesting that common-cold HCoV cross-reactivity is variable and likely to be conditioned by HLA. It is plausible that patients carrying these HLAs may exhibit more robust protection against SARS-CoV-2 and its variants. We lastly identified a set of 10 peptides that are highly conserved across multiple coronavirus strains, to serve not only as potential pan-coronavirus T cell targets, but we propose are leading candidates as cross-reactive CD8+ T cell epitopes.

## Results

### Curation of functionally evaluated SARS-CoV-2 peptides

To investigate the potential for T cell cross-reactivity against SARS-CoV-2 as conferred by common-cold HCoVs, we curated a comprehensive pool of SARS-CoV-2 class I and II peptides from three datasets (see Methods), which have been functionally evaluated for CD4+ and CD8+ T cell responses (see Figure 1 for study overview). The data comprise 1799 and 1005 immunogenic and non-immunogenic SARS-CoV-2 *peptides* respectively (Fig 2A). Many of these peptides were tested for T cell reactivity in the context of multiple HLA alleles and/or by multiple assays (IFNγ, IL-5 production etc). Furthermore, some peptides are described by qualitative labels corresponding to varying response magnitude (Positive-high and Positive-low etc). Taking these combinations into account, we found 3979 and 2427 immunogenic and non-immunogenic *complexes* (Fig 2B). For immunogenic complexes, the most common lengths are 9-mers, followed by 15- and 10-mers (Fig 2C), and 36.0% are presented by class I MHC, 32.9% by class II (Fig 2D) and for 31% MHC type is unknown (Fig S1A). For non-immunogenic complexes, 36.1% are presented by class I, 26.4% by class II and for 37.51% the MHC is unknown. At the gene level, HLA-allele specific information was available for 934 (56.5%) and 607 (42.2%) of immunogenic class I and II complexes respectively (Fig S1A).

**Figure 1:**
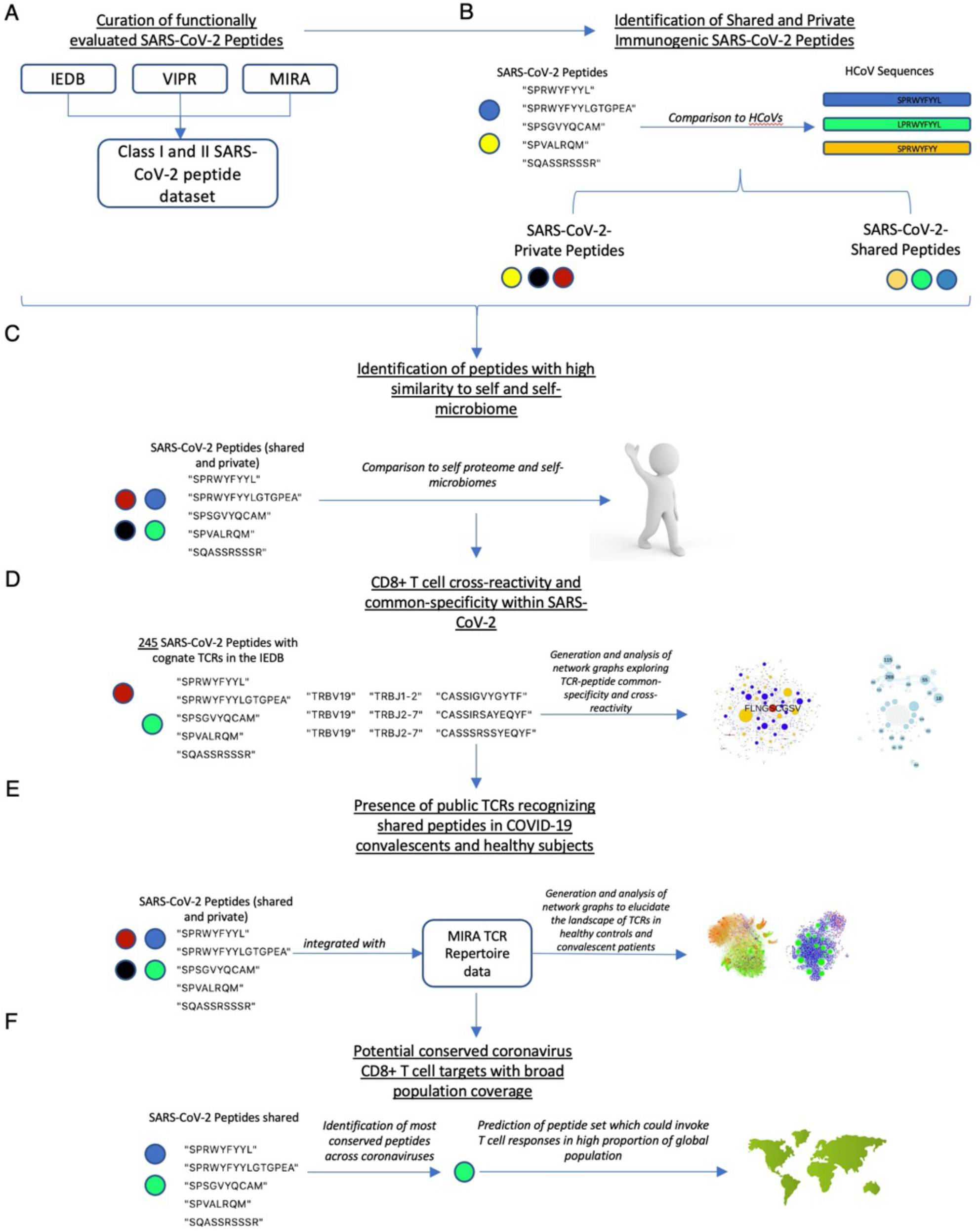
Overview of the study. A) Functionally evaluated SARS-CoV-2 peptides are gathered from three online repositories. Data are cleaned and integrated, thus curating a comprehensive pool of SARS-CoV-2 class I and II peptides. Exploratory data analysis followed. B) Next, each immunogenic SARS-CoV-2 peptide of length l is compared to each possible linear peptide of length l from four common-cold causing human coronavirus strains. Based on similarity criteria and following confirmation that the target hit from HCoV is predicted to bind HLA, peptides are classified as ‘shared’ SARS-CoV-2-HCoV peptides. Those which do not adhere to the criteria are classified as SARS-CoV-2 private. C) The entire set of SARS-CoV-2 shared and private, immunogenic and non-immunogenic peptides is compared to the human proteome and gut and airways microbiomes. D) 245 peptides from our SARS-CoV-2 peptide dataset have known cognate TCRs in the IEDB. These peptide-TCR associations were examined to explore the extent of cross-reactivity and common-specificity within SARS-CoV-2. E) Both shared and private immunogenic SARS-CoV-2 peptides are integrated with the COVID-19 MIRA TCR repertoire dataset and employed to examine the presence of TCRs recognizing shared/private SARS-CoV-2 peptides in health and/or disease. F) The entire set of 126 shared SARS-CoV-2 peptides is searched for those most highly conserved across coronaviruses. This resulted in 17 peptides, of which we used to predict global and regional population coverage, given predicted HLA alleles.

**Figure 2:**
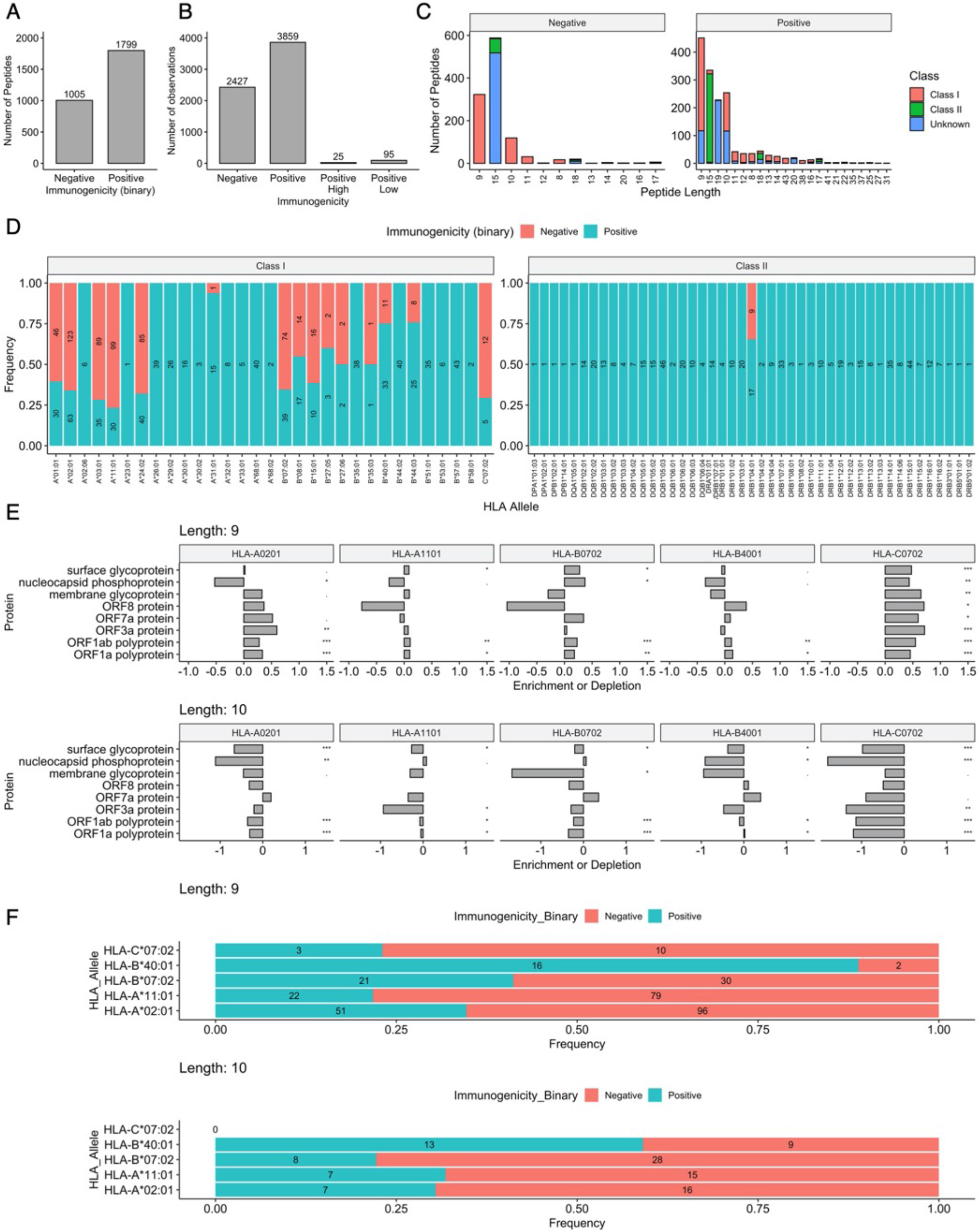
A comprehensive pool of functionally validated SARS-CoV-2 peptides. Barplots showing A) The number of SARS-CoV-2 peptides deemed ‘positive’ or ‘negative’. A ‘negative’ label reflects that for a peptide, there are only negative (nonimmunogenic) qualitative observations while ‘positive’ reflects at least one immunogenic observation. B) The number of total pMHC complexes observed in the dataset, including all assay and HLA combinations for each peptide. C) The distribution of lengths of complexes amongst our SARS-CoV-2 dataset. Left plot shows nonimmunogenic ‘Negative’ peptides. Right plot shows immunogenic or ‘Positive’ peptides. MHC class is colour coded. D) The frequencies of immunogenic or non-immunogenic class I or II observations, for peptides where specific HLA allele information is available. Numeric labels show the number of peptides in each group. E) The logs odd ratio of observed and expected number of presented peptides of lengths 9 and 10 by common HLA alleles for SARS-CoV-2 proteins > 100 amino acids in length. Significance calculated using binomial distribution. F) The frequency of immunogenic and nonimmunogenic peptides as presented by common class I HLA alleles arising from SARS-CoV-2. Numeric labels show the number of observations per immunogenicity status.

Given the high proportion of missing MHC information, we employed netMHCpan 4.1 and netMHCIIpan to predict presenting class I and class II alleles respectively for immunogenic peptides (see Methods). Here, we were able to identify 98% of known MHC molecules, providing confidence in predictions for unknown alleles (Fig S1B).

We next sought to examine whether HLAs exhibit preferences towards presenting peptides from certain SARS-CoV-2 proteins. By employing a similar methodology to Karnaukhov et al^17^, we gauged the enrichment and depletion of HLA ligands arising from these proteins (see Methods). Indeed, we observed differential antigen presentation by HLAs e.g., HLA-C*07:02 appears to be the most consistently enriched in presenting 9mers from the examined proteins (Fig 2E), while HLA-A*02:01 is enriched in presenting 9mers from ORFs but depleted for 10mers across most assessed proteins. This disparity may be due to a known preference of 9-mers for HLA-A*02:01 ^18^. Furthermore, despite the prevalence of HLA A*02:01 in the global population and in MHC presentation experiments, this allele appears to be depleted for presenting ligands from SARS-CoV-2 proteins that have been the focus of intense experimental work e.g., spike and nucleocapsid phosphoprotein.

These patterns of HLA preferences in presenting SARS-CoV-2 peptides appear to differ for 9- and 10-mers. For example, whereas HLA-C*07:02 is enriched for presenting 9mers, this allele appears to be a poor presenter of 10mers from each examined protein. It is unclear why substantially fewer 10-mer HLA-C*07:02 ligands are predicted than 9mers, however it is plausible that this allele may prefer 9mers, as appears to be the case with HLA-A*02:01, - A*11:01 and -B*40:01 ^18^, or that this may be a SARS-CoV-2 specific effect.

Although it is of great interest to reveal the rate to which SARS-CoV-2 MHC-bound peptides are immunogenic in humans ^19^, it cannot be examined directly with existing data because not all MHC-bound SARS-CoV-2 peptides have been evaluated for immunogenicity. Nevertheless, we explored the pool of MHC-bound peptides in our dataset that have been examined for a T cell response, to gauge the proportion that SARS-CoV-2 pMHC are immunogenic. Overall, we observed low rates of immunogenic pMHC (Fig 2F), although ligands of HLA-B*40:01 appear to be commonly immunogenic. Interestingly we observed that HLA-C*07:02 does not present any 10-mers in our dataset. This apparent preference for 9-mers is consistent with availability of HLA-C*07:02 ligands tested for T cell response in humans from the IEDB, where there exist only 121 unique peptides, of which 73% are 9mers and only 12% are 10mers. In summary, these data suggest length and source protein preferences for HLA alleles presenting SARS-CoV-2 peptides and that HLA-B*40:01 SARS-CoV-2 ligands are commonly immunogenic.

### Identification of Shared and Private Immunogenic SARS-CoV-2 peptides

To discriminate SARS-CoV-2-HCoV shared (herby referred to as ‘sCoV-2-HCoV’) peptides, we compared immunogenic SARS-CoV-2 peptides to HCoV protein sequences. For this, we define a metric that considers 1) sequence homology, 2) physicochemical similarities (MatchScore^20^) and 3) presentation status for which the source peptide from SARS-CoV-2 and the target peptide from one of the HCoVs are required to be presented by the same HLA. A source peptide is defined as shared if it fulfils all these three conditions otherwise is considered as a private peptide (see Methods).

Using our metric, we identified 126 unique SARS-CoV-2 (immunogenic) peptides pointing to 285 highly similar peptides in HCoVs (Supplementary Data File 1). Hence, we provide a comprehensive map of private and shared SARS-CoV-2 functionally evaluated immunogenic peptides, and for sCoV-2-HCoV peptides, their matches from each HCoV.

Out of the HLAs tested (see Methods) 33 and 28 class I and II HLAs respectively were predicted to present the target HCoV pMHCs (Fig3A). HLA-A*02:01 and HLA-B*27:05 were the most and least common class I presenters respectively. For class II, DRB1-1501 and DRB5-0101 were the most common presenters, while DRB1-0301 and DRB1-1303 were the least. Most shared class I and II peptides were predicted to bind multiple HLA allelic variants (Fig 2SA). Compared with private peptides it appears that sCoV-2-HCoV peptides are presented by less HLAs, although this was not significant (Fig S2C). Nevertheless, the range of predicted alleles for these peptides suggests recognition in broad geographical and ethnic settings ^21^.

**Figure 3:**
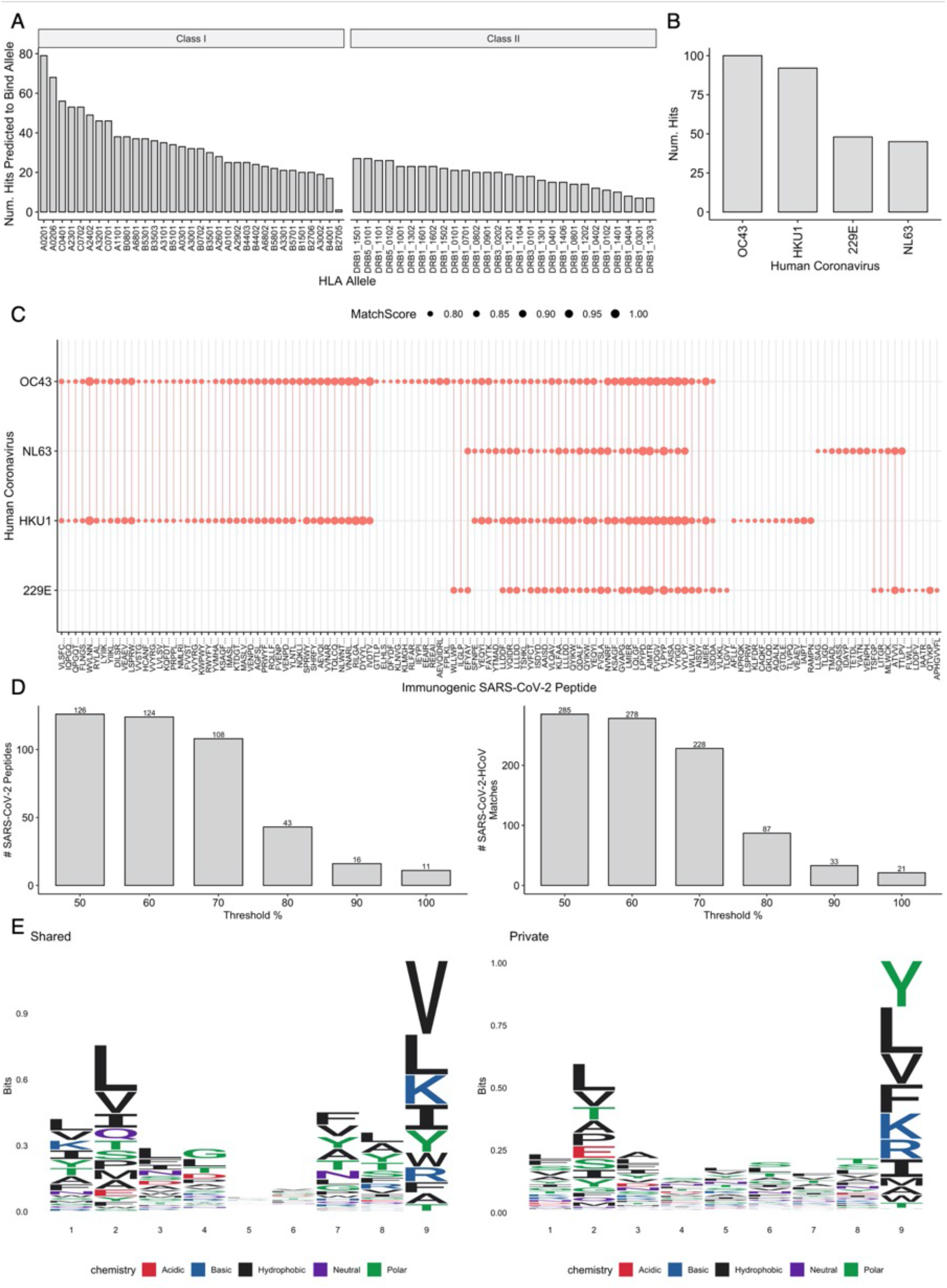
A set of peptides from human coronavirus strains with high similarity to immunogenic SARS-CoV-2 peptides. A) A barplot showing the number of high similarity matches predicted to bind a set of common HLA class I and class II alleles. B) A barplot showing the number of unique high similarity matches derived from each human common-cold-causing coronavirus. Each hit is defined as a unique observation with MatchScore > 0.75, between an immunogenic SARS-CoV-2 peptide with length x, and a stretch of length x from one viral protein. C) A dot and line plot showing each SARS-CoV-2 peptide and to which common-cold-causing coronavirus it exhibits a high similarity match. The size of each point reflects the MatchScore, i.e the primary similarity metric. D) Barplots showing the number of unique SARS-CoV-2 peptides (left) and SARS-CoV-2-HCoV matches (right), at different thresholds of the sequence homology metric, i.e the % of the amino acids that must be conserved between the SARS-CoV-2 peptide and its HCoV match. E) Sequence logo plots comparing amino acid usage amongst SARS-CoV-2 shared and private 9-mer peptide sequences.

For the 126 SARS-CoV-2 peptides with high similarity to HCoV, we also observed binding to multiple HLAs (FigS2B). In addition, we found that 9mers comprise 54% of the 126 SARS-CoV-2 peptides with high-similarity matches to HCoV, followed by 15mers (19%) and 10mers (17.5%) (FigS2C). Consistent with previous reports ^22^, the betacoronaviruses HKU1 and OC43 were most enriched in target matches (Fig3B), perhaps due to higher total sequence homology among betacoronavirus strains ^23^. We next examined the extent to which immunogenic SARS-CoV-2 peptides exhibit homology to *multiple* HCoV strains. Surprisingly, we found that 42 SARS-CoV-2 immunogenic peptides exhibit matches to at least three strains (Fig3C). However, we observed small clusters of peptides that only possess homology with one strain, e.g OC43 or HKU1. ORF1ab protein and spike surface glycoprotein produced the highest quantity of shared SARS-CoV-2-HCoV peptides in both strains, and the protein regions from which these peptides were found are similar in both HKU1 and OC43 (Fig S2E-H).

Of particular note about our map of shared and private peptides is that this map is subject to thresholds that we used in our metric. The sequence homology threshold that was used here is 50% and most peptides had greater than or equal to 70% sequence homology (Fig S2E) Although, more stringent sequence homology parameter will result a map containing fewer shared peptides (Fig 3D), our main conclusions in this manuscript remain the same even with sequence homology threshold of 70% (data are not shown).

Lastly, we compared the amino acid distribution between shared and private SARS-CoV-2 peptides for 9-mers, which is the most common peptide length in our dataset (Fig 3E). We observed some moderate differences, e.g., increased prominence of Valine at position 9 within shared peptides.

We have therefore identified a pool of 126 SARS-CoV-2 immunogenic peptides - that exhibit high similarity to 291 peptides in HCoV strains - which are likely to be presented by an array of class I and II HLA molecules. This array of presenting alleles suggests the potential for broad global population coverage, which is explored later. We propose that this pool of experimentally confirmed immunogenic SARS-CoV-2 peptides and their counterpart high similarity matches be considered as potential targets for T cell cross-reactivity, therefore warranting investigation into pre-existing immune memory from HCoV or a role in protection from SARS-CoV-2 variants.

### Identification of peptides with high similarity to self and self-microbiomes

To prevent aberrant T cell mediated inflammation and tissue damage, the immune system has evolved several checkpoint mechanisms. These include thymus negative selection and peripheral tolerance. Indeed, dissimilarity to self is increasingly recognised as a component of peptide immunogenicity ^25^, which may assist in calibrating a balance between immunogenicity and inflammatory pathogenesis.

To evaluate the extent to which dissimilarity to self and self-microbiomes contribute to SARS-CoV-2 peptide immunogenicity, we took a similar approach and used our metric to compare SARS-CoV-2 peptides to human self-proteome and microbiomes that include 457 gut and 50 airway microbiota. (see Methods). Here, for SARS-CoV-2 HLA class I presented 9- and 10-mer peptides we observed that immunogenic SARS-CoV-2 peptides were significantly more dissimilar to the human proteome than their non-immunogenic counterparts (Fig 4A, S3A). Using this approach, we could not detect any significant difference between immunogenic and none immunogenic class II peptides in their dissimilarity to self-proteome (Fig S3B).

**Figure 4:**
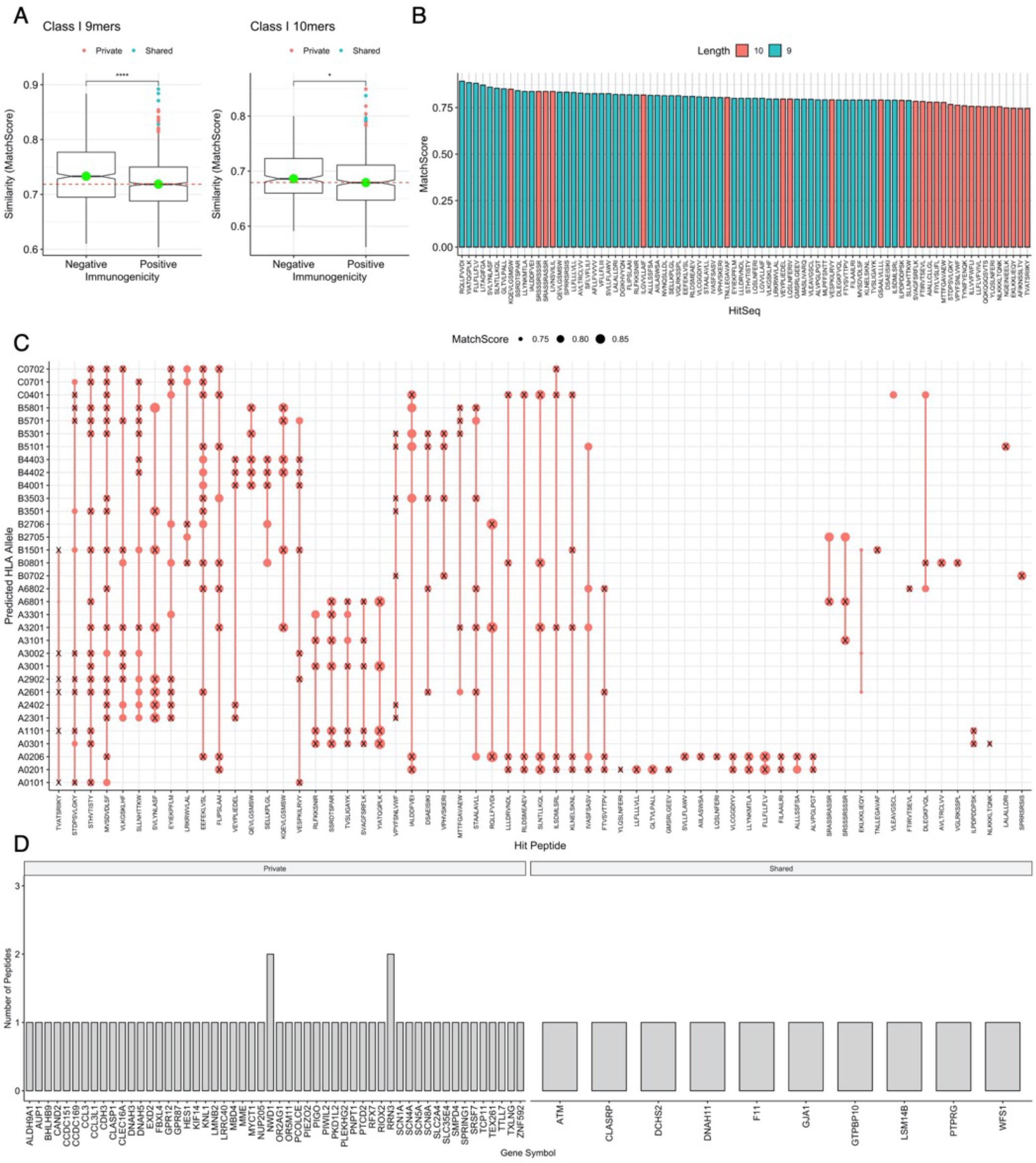
A pool of immunogenic SARS-CoV-2 peptides with high similarity to human genes. A) Notched boxplots showing the similarity (evaluated by the MatchScore) of nonimmunogenic and immunogenic SARS-CoV-2 peptides with sequences derived from the human proteome of lengths 9 (immunogenic n = 906, nonimmunogenic n=734 peptides) and 10 (immunogenic n=394 and nonimmunogenic n=394 peptides). The green dot shows the median of each group, red line represents the median of the immunogenic group. B) A barplot showing the MatchScores of hits with high similarity to the human proteome, colour-coded by protein length. C) A dotplot showing the predicted HLAs of candidate peptides with high similarity to SARS-CoV-2 proteome. The size of the point reflects the MatchScore. An ‘X’ shows where the SARS-CoV-2 derived peptide and the match are predicted to bind the same allele. D) A bar chart showing the genes from which the high similarity match peptides arise in the human proteome, separated by the “Private” or “Shared” epitope status of the SARS-CoV-2 epitope.

Interestingly however, for peptides of both lengths 9 and 10, we identified several immunogenic SARS-CoV-2 peptides with considerable sequence similarity to the human proteome (Fig 4A-B, Table S1). For the top 10% of these peptides with highest similarity to self, the mean amino acid conservation (the proportion of the amino acid sequence which is exactly conserved) between these peptides and corresponding self matches is 72.1% with 8.33% standard deviation (see Supplementary Data File 2 for number of substitutions under column ‘Hamming’). In general, T cells specific for these peptides should be subject to negative selection otherwise it is plausible that aberrant immune responses may occur during the course of the disease in the form of immunopathology or in the future in the form of autoimmunity ^8,26,27^.

To investigate the potential association of these peptides in immunopathology further, we predicted MHC presentation by a set of class I HLA alleles (see Methods) for the top 10% of peptides most similar to the human proteome for 9mers and 10mers. We observed that these peptides with high similarity to self are predicted to bind multiple HLAs (Fig 4C), and interestingly, we found that in most cases, the SARS-CoV-2 immunogenic peptide and the match from the human proteome are predicted to be presented by the same allele (Fig4C).

Next, we examined the list of genes with high sequence similarities to these SARS-CoV-2 immunogenic peptides (Table S1 & Supplementary Data File 2). Of particular interest, we found e.g. CCL3 and CCL3L1 which are linked to cytokine storms and the expression of which have been reported to be elevated in severe COVID-19 patients ^28–33^ (Fig 4D, Supplementary Data File: 1). We additionally observed CD163, similarly associated with severe COVID-19, however the predicted presentation score of HLA-B*15:01 for peptides from CD163 with high similarity to SARS-CoV-2 were slightly beyond the generally accepted ‘binding’ cutoff. Interestingly, the SARS-CoV-2 peptides exhibiting sequence homology to CCL3 and CCL3L1 (and CD163) were private to SARS-CoV-2 (Fig 4D & Table 1) - which may increase the likelihood of being involved in immunopathology after infection. Additionally, we observed considerable amino acid conservation with matches from these genes, with 77.8% for 9mers and 70% for 10mers (Table 1).

**Table 1:** SARS-CoV-2 peptides with high similarity to the human proteome, from genes reported to be involved with severe COVID-19. Note, peptides derived from CD163 were not predicted to bind HLA.

| Human Match | MatchScore | SARS-CoV-2 Peptide | HLA Allele | Shared/Private | Protein | Length | Proportion Conserved |
| --- | --- | --- | --- | --- | --- | --- | --- |
| STAALAVLL | 0.806 | STAALGVLM | HLA-A*26:01 | Private | >sp\|P10147\|CCL3_HUMAN C-C motif chemokine 3 OS=Homo sapiens OX=9606 GN=CCL3 PE=1 SV=1 | 9 | 0.778 |
| STAALAVLL | 0.806 | STAALGVLM | HLA-A*26:01 | Private | >sp\|P16619\|CL3L1_HUMAN C-C motif chemokine 3-like 1 OS=Homo sapiens OX=9606 GN=CCL3L1 PE=1 SV=1 | 9 | 0.778 |
| ILGVVLLAIF | 0.818 | IVGVALLAVF | HLA class I | Private | >sp\|Q86VB7\|C163A_HUMAN Scavenger receptor cysteine-rich type 1 protein M130 OS=Homo sapiens OX=9606 GN=CD163 PE=1 SV=2 | 10 | 0.7 |
| ILGVVLLAIF | 0.818 | IVGVALLAVF | HLA class I | Private | >tr\|F5GZZ9\|F5GZZ9_HUMAN Scavenger receptor cysteine-rich type 1 protein M130 OS=Homo sapiens OX=9606 GN=CD163 PE=1 SV=1 | 10 | 0.7 |
| ILGVVLLAIF | 0.818 | IVGVALLAVF | HLA class I | Private | >tr\|H0YFM0\|H0YFM0_HUMAN Scavenger receptor cysteine-rich type 1 protein M130 (Fragment) OS=Homo sapiens OX=9606 GN=CD163 PE=1 SV=1 | 10 | 0.7 |
| ILGVVLLAIF | 0.818 | IVGVALLAVF | HLA class I | Private | >tr\|C9JHR8\|C9JHR8_HUMAN Scavenger receptor cysteine-rich type 1 protein M130 OS=Homo sapiens OX=9606 GN=CD163 PE=1 SV=1 | 10 | 0.7 |

CCL3 and CCL3L1 are both ligands for CCR1 and CCR5. Interestingly, CCR1 variants are linked to pulmonary macrophage infiltration in severe COVID-19^34^ and inhibition of CCR5 in critical COVID-19 patients has been associated with a decrease in plasma IL-6 and SARS-CoV-2 RNA and an increase in CD8+ T cells ^35^. Additionally, intermediate monocytes which constitutively express high levels of CCR5 have recently been suggested as playing a role in post-acute sequelae of COVID-19 ^36^ (often referred to as ‘long-COVID’). Of further interest, we found SMPD4 and SLC1A4, which together with CCL3 and CCL3L1 are involved in the response to TNF, which is part of the cytokine storm following COVID-19 disease.

By comparing SARS-CoV-2 peptides to human microbiomes, we observed subtle higher dissimilarity of SARS-CoV-2 immunogenic peptides to the gut (Fig S3C) and airways (Fig S3D) microbiomes, which may suggest a link between the diversity of both microbiota and heterogeneity of the disease in populations, although this warrants further investigation.

Given the magnitude of the global pandemic and the widespread vaccination required to combat it, future virus-induced autoimmune disease and immunopathology is of concern. Overall, this analysis suggests dissimilarity of viral peptides to self-proteins as a correlate of peptide immunogenicity. Furthermore, we present candidate genes and peptides with high similarity to SARS-CoV-2 T cell targets, which we suggest as prime targets for further investigations into their role in autoimmune disease and immunopathology following SARS-CoV-2 infection and/or vaccination.

### CD8+ T cell cross-reactivity and common-specificity within SARS-CoV-2

A valuable characteristic of our map of SARS-CoV-2 shared and private peptides, is that for 245 of these (out of 1279 class I immunogenic peptides), cognate TCRs at the beta chain resolution are available in the IEDB. We therefore set out to map the TCR landscape through a network approach to explore the potential for cross-reactivity among SARS-CoV-2 specific CD8+ T cells, and their common-specificity. Here, to avoid overestimating connectivity, any peptides of different lengths, which share starting positions in the SARS-CoV-2 proteome and are recognised by identical sets of TCRs, are considered as one peptide.

Through a two-mode (bipartite) network-graph illustrating the connectivity of SARS-CoV-2 immunogenic peptides with their cognate TCRs, amongst a highly connected topography we observed considerable connectivity for some sCoV-2-HCoV peptides e.g. “FLN..” (Fig S4A). Exploring this further, we projected the bipartite network-graph into a one-mode graph where nodes represent peptides and an edge between two nodes requires existence of a TCR recognising both peptides (Fig S4B). The clustering around a small set of hubs suggests that many experimentally assessed TCRs target a small set of SARS-CoV-2 peptides. Indeed, we found that in this dataset, 80% of the TCRs are reported to recognise only 40 (16%) peptides, of which 4 are sCoV-2-HCoV shared peptides and 36 are SARS-CoV-2 private (Fig S4C). This dominant set of peptides may be due to experimental biases e.g., research may be heavily biased toward several protein regions. However, this may also reflect a selection bias by SARS-CoV-2 specific TCRs. In this regard, amongst these 80% of TCRs, we observed high usage of V gene TRBV20-1 ^37^ and J gene TRBJ2-1 ^38^ (Fig S4D), that have been previously reported to have implications in COVID-19 patients.

Similarly, we examined the extent of common specificity in SARS-CoV-2 specific T cells by a one-mode graph in which nodes represent TCRs and an edge represents whether two nodes (TCRs) recognize the same peptide (Fig S4E). Interestingly, this graph reveals a set of highly connected hubs reflecting levels of common specificity, however there are many TCRs which recognise only a single unique peptide. Comparing these two sets of TCRs, we did not observe considerable differences in their CDR3β sequences (Fig S4F-G), however we observed differences in V and V-J gene usage (Fig S4H-J).

In summary, we employed peptides with known cognate TCRs in the IEDB database - although limited in numbers – to explore SARS-CoV-2 CD8+ T cell cross-reactivity. Our network approach demonstrates that SARS-CoV-2 CD8+ T cells can cross-react and exhibit common-specificities.

### Presence of public TCRs recognising sCoV-2-HCoV peptides in COVID-19 convalescents and healthy subjects

We next integrated our map of SARS-CoV-2 shared and private peptides with a recently published dataset known as ‘MIRA’ ^39^ to track the patterns of public TCRs recognizing sCoV-2-HCoV peptides in convalescents and/or healthy subjects. Here, Nolan et al., employed the ‘Multiplex Identification of Antigen-Specific T cell receptors’ (MIRA) assay to identify SARS-CoV-2 specific TCRs from PBMCs and naïve T cells. These data include more than 160k high confidence SARS-CoV-2-specific TCRs mapped to target peptides from 39 Healthy controls (HC) (defined as unexposed to SARS-CoV-2) and 90 COVID-19 convalescent patients. These data consist of 792 unique SARS-CoV-2 peptides, 54 of which are sCoV-2-HCoV shared peptides.

To elucidate the landscape of public TCRs in HC and convalescent patients, we generated a bipartite graph comprising all public TCRs (defined as CDR3b+V +J gene(s) present in at least two subjects) cognate for SARS-CoV-2 private and sCoV-2-HCoV shared peptides (Fig 5A). This graph revealed two clear hubs. In the first (green nodes), we observed that healthy subjects were connected to public TCRs which recognise both sCoV-2-HCoV and SARS-CoV-2-private peptides. In the second hub (red nodes) comprising convalescent patients, we observed that generally their public TCR repertoires predominately recognise SARS-CoV-2-private peptides. Indeed, it appears that cognate TCRs of sCoV-2-HCoV peptides are more pronounced in HC (Fig S5A-Shared, wilcoxon p=0.00029) whereas cognate TCRs of SARS-CoV-2-private peptides appear enriched in the convalescent cluster (Fig S5A-Private). Interestingly, we observed a considerable number of TCRs recognising shared peptides which are common between these two subject clusters, indicating that sCoV-2-HCoV-specific public TCRs are present not only in COVID-19 patients but are also expanded from unexposed individuals (Fig 5A-B).

**Figure 5:**
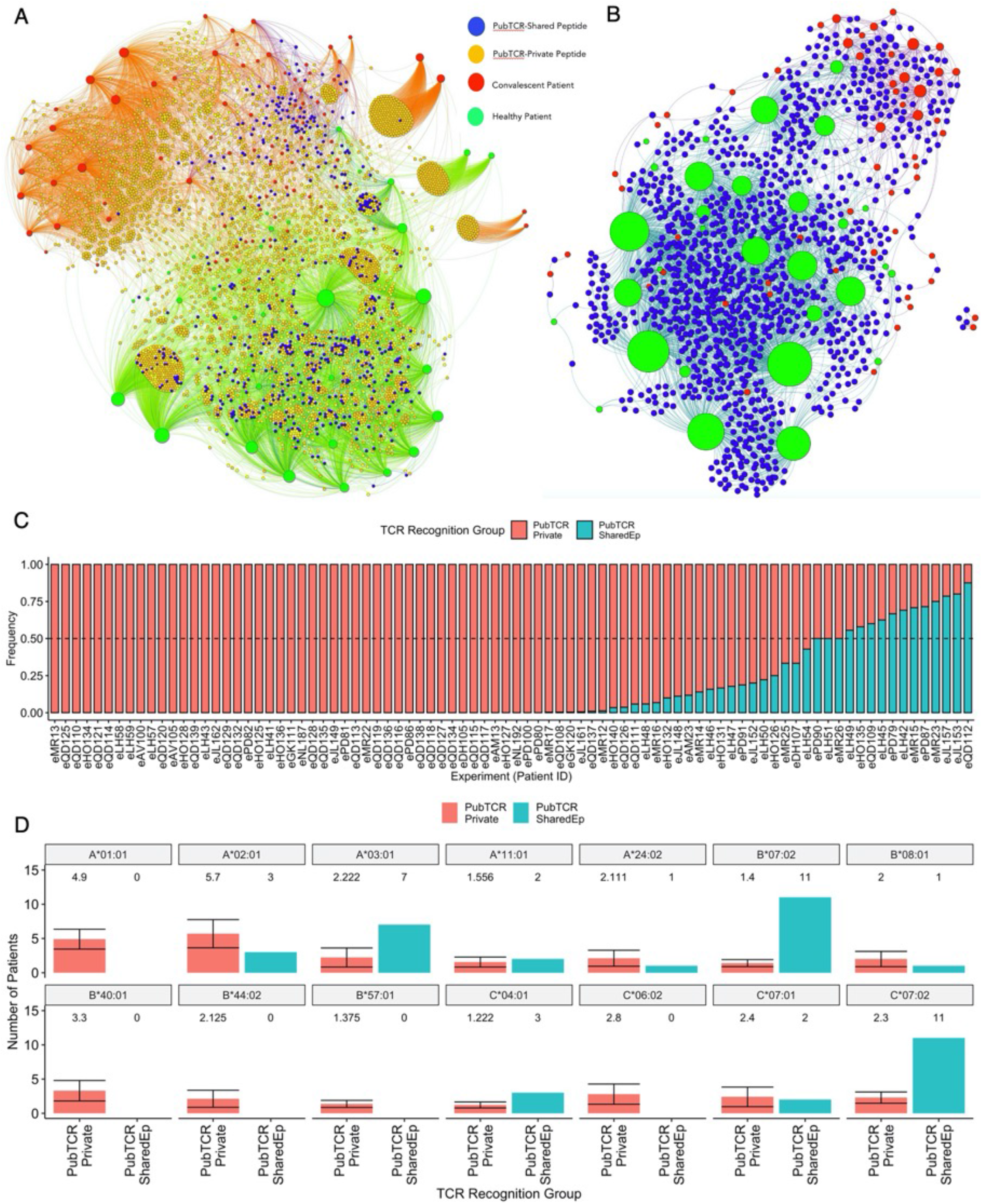
A landscape of T cell responses against SARS-CoV-2-HCoV Shared and SARS-CoV-2-private peptides in healthy or COVID-19 convalescent individuals: A) A bipartite network graph showing SARS-CoV-2-specific public TCRs which recognize shared or private SARS-CoV-2 peptides in healthy or convalescent patients. TCRs are colour-coded by whether they recognize only shared peptides (blue), only private peptides (orange) or both (yellow). COVID-19 convalescent patients are labelled red while healthy controls are labelled green. Node size reflects degree of connectivity, i.e. the quantity of an individual’s TCRs which are shared with other patients. B) A bipartite network graph showing SARS-CoV-2 public TCRs that recognize SARS-CoV-2-HCoV shared peptides. Patient node size reflects the quantity of their TCRs which are shared with another patient. Healthy patients are labelled green, COVID-19 convalescent are labelled red, and (public) TCRs are labelled blue. C) A barplot showing the frequency that each convalescent patient’s public TCRs recognize SARS-CoV-2-private (red) or SARS-CoV-2-HCoV-shared (blue) peptides. Patients with identical frequencies are ordered by the number of TCRs. D) Barplots showing the quantities of COVID-19 convalescent patients who carry 14 class I HLA alleles of interest. Patients are grouped by whether their public TCRs predominately recognize “Private” (PubTCR_Private, n=12, sampled 10 times) or “Shared” (PubTCR_SharedEp, n=12) peptides. For the PubTCR-Private group, 12 patients were sampled 10 times and the number of patients carrying alleles was measured. The mean number of patients carrying each allele and the error are visualized. For the PubTCR-SharedEp group, the data contain only 12 patients of interest, thus the number of patients carrying each allele is measured and visualised.

Given that in these healthy donors, the TCRs are generally from naïve CD8+ T cells which are expanded and stimulated with SARS-CoV-2 peptide pools and analysed with the ‘MIRA’ assay, the presence of cognate TCRs recognising sCoV-2-HCoV peptides in HC as well as COVD-19 patients may not necessarily translate into pre-existing T cell immunity. Rather, due to the high similarity between the cognate SARS-CoV-2 antigens and (predicted) HCoV presented peptides, we suggest it is plausible that these SARS-CoV-2 specific TCRs are cross-reactive with HCoV peptides. Indeed, consistent with Francis et al.,^15^ who demonstrate pre-existing memory CD8+ T cells to SPR* peptide in 80% of unexposed individuals, we found a set of public TCRs - which are observed in both convalescent and unexposed individuals - recognising this sCoV-2-HCoV peptide. In this light, we reveal candidate public TCRs and corresponding SARS-CoV-2 peptides with high similarity to HCoVs, which should be examined further for cross-reactive potential.

From these two bipartite graphs, we observed that healthy individuals respond to a balance of SARS-CoV-2 private and sCoV-2-HCoV peptides, although it appears that infection *primarily* dictates a dominant recognition of private SARS-CoV-2 peptides (Fig S5B). For convalescent patients, we observed that public TCR repertoires of the majority (51/86) of patients are almost entirely (>=99%) occupied by TCRs recognising SARS-CoV-2 private peptides (Fig 5C). However, in a subset of convalescent patients, public TCRs recognising sCoV-2-HCoV peptides comprise a substantial fraction of the public repertoire. In fact, for 12 convalescent patients, >50% of their public TCRs recognise sCoV-2-HCoV peptides.

Comparing these two groups of patients, we did not find evidence of a link toward biological sex or age. To explore potential correlates, we first gathered the 12 patients whose public TCRs most dominantly (>50%) recognise sCoV-2-HCoV peptides (labelled *PubTCR-SharedEp*) and then via sampling 12 patients 10 times from the set of 51 patients whose public TCRs almost entirely recognise SARS-CoV-2 private peptides (labelled *PubTCR-Private*), we compared HLA coding genes of these two groups. We observed that the *PubTCR-SharedEp* group is statistically enriched for carrying HLA-B*07:02, HLA-C*07:02 and HLA A*03:01, whereas the former group includes a broader set of HLAs among which HLA A*01:01 was more pronounced (Fig 5D). The enrichment of HLA-B*07:02 in the *PubTCR-Shared*Ep group is consistent with recent work from Francis et al^15^, and these data are in agreement with their claim that CD8+ T Cell HCoV-CoV-2 cross-reactivity may be conditioned by HLA.

Employing these two groups and sampling a set of healthy patients (n=12), we reveal the set of epitopes only recognised by public TCRs in these healthy patients, and those shared with the convalescent *PubTCR-SharedEp* group (Fig S5C, Supplementary Data File 3 and 4). Additionally, we reveal the peptides only observed in the *PubTCR-Private* convalescent group, adding to previous insights that SARS-CoV-2 infection can provoke T cell responses to a novel set of peptides compared to those expanded from unexposed patients ^7^.

Recent work shows cross-reactive *private* TCRs from unexposed subject repertoires, capable of recognising both the SARS-CoV-2 SPR* peptide and its <u>L</u>PR* homolog from HCoVs OC43 and HKU1. By mapping out which SARS-CoV-2 peptides are recognised in which individuals by private TCRs, we observed SPR* but also an additional set of sCoV-2-HCoV peptides recognised in both healthy and convalescent patients (Fig S5D, Supplementary Data Files 5-6). Lineburg et al., ^40^ recently reported private TCRs in HLA-B*07:02^+^ unexposed individuals which cross-react with both the SARS-CoV-2 SPR* peptide and the OC43/HKU1 homolog <u>L</u>PR*, which indicates a level of pre-existing immunity. Of these TCRs, we found two (defined as CDR3b, TRBV, TRBJ) which appear in two HLA-B*07:02^+^ unexposed individuals within the MIRA dataset (Table S2). As these TCRs are now observed in two separate datasets, we therefore propose these as public TCRs, capable - as identified by Lineburg et al., - of cross-reacting with both SARS-CoV-2 SPR* and OC43/HKU1 LPR* peptides.

Taken together, we report existence of a set of CD8+ TCRs in both HC and COVID-19 convalescent patients that recognise SARS-CoV-2 peptides with high sequence similarity to a pool of predicted HCoV pMHC. This high sequence similarity indicates cross-reactive potential of these TCRs. Primarily however, we observed that COVID-19 patients develop public TCR responses to private peptides – many of which are not observed in unexposed individuals - indicating that any cross-reactive potential is limited. For the subset of COVID-19 patients whose public TCRs are directed towards sCoV-2-HCoV peptides - and are observed in HC - we found distinct HLA profiles. Therefore, in agreement with recent data from Francis et al., we suggest that CD8+ T cell HCoV-CoV-2 cross-reactive potential is apparent, although likely conditioned by patient HLA genotype. It is plausible that these patients may exhibit more robust protection against SARS-CoV-2 and its variants.

### Potential conserved coronavirus CD8+ T cell targets with broad population coverage

Given the emergence of new SARS-CoV-2 variants and concern over the theoretical capacity of future mutants to evade current vaccine strategies^1^, conserved CD8+ T cell targets across multiple coronavirus strains with the potential to elicit T cell responses in a large percentage of global populations are of interest. We therefore searched our peptide map for SARS-CoV-2 peptides with ‘high-similarity’ matches to *multiple* HCoVs, and with cognate TCRs in the MIRA dataset. To select only the top ‘high-similarity’ SARS-CoV-2-HCoV matches for this analysis, we applied a more stringent sequence homology threshold. Indeed, in addition to the ‘MatchScore’ and peptide presentation criteria outlined previously (see Methods: Discriminating shared and private SARS-CoV-2 peptides), we only retained matches with at least 70% sequence conservation (i.e. allowing 30% amino acid substitution).

We found 86 peptides that match these criteria, 84 of which are recognised by TCRs in both convalescent and HC (Fig 6A-B). We next focused on SARS-CoV-2 peptides with high similarity matches in >=3 HCoV strains (Table 2, Supplementary Data File 7). Of these SARS-CoV-2-HCoV matches, the number of amino acid substitutions ranged between 0-3, with a mean of 1.79 and standard deviation of 0.78. Additionally, while each of these peptides exhibited a high similarity match to either MERS or SARS-CoV, the majority exhibited homology with both of these viruses (Fig S6A). As well as high conservation across many coronavirus strains, collectively these SARS-CoV-2 peptides are predicted to bind multiple HLA alleles (Fig 6C), raising the possibility that this set of peptides may elicit T cell responses in a substantial proportion of the global population.

**Figure 6:**
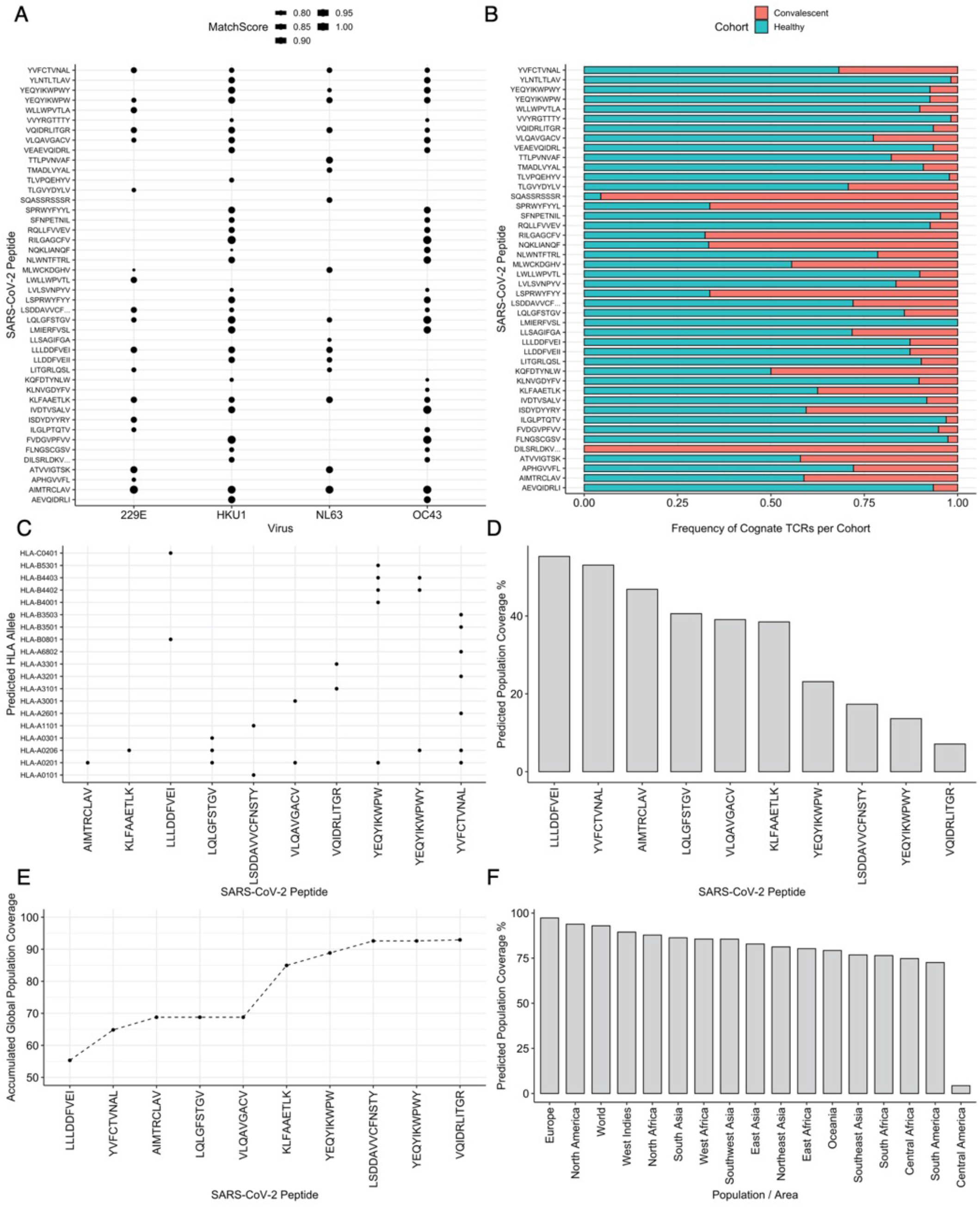
T cell epitopes with known cognate TCRs which are conserved across multiple HCoVs and SARS-CoV-2 exhibit broad population coverage: A) A dot plot showing SARS-CoV-2 peptides with high similarity to more than one HCoV that are recognized by TCRs in the MIRA dataset. Size of the dot represents the MatchScore. B) The frequency of cognate TCRs which recognize these peptides from the COVID-19 convalescent or healthy cohorts. C) The HLA alleles predicted to present SARS-CoV-2 peptides with high similarity matches to 3 or 4 HCoV strains. D) Global population coverage as calculated by the ‘IEDB population coverage tool’ for each individual SARS-CoV-2 peptide with high similarity matches to 3 or 4 HCoV strains. E) Accumulated global population coverage predicted by the IEDB population coverage tool. F) Regional population coverage for the entire set of 10 SARS-CoV-2 peptides with matches to 3 or 4 HCoV.

**Table 2:**
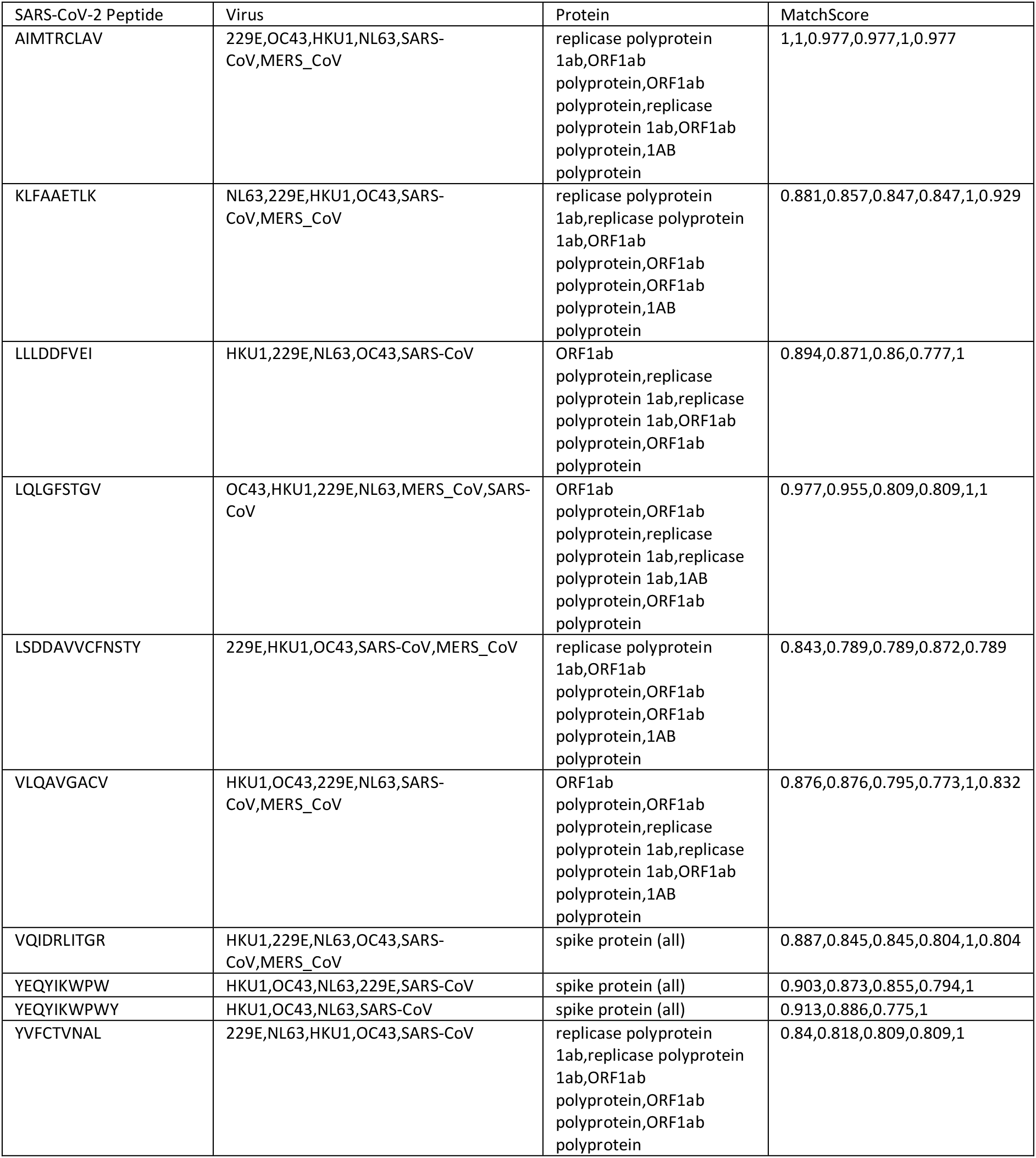
Highly conserved CD8+ T cell peptides across SARS-CoV-2 and HCoV strains, with high population coverage

We next sought to determine the extent in global and regional populations that these CD8+ T cell targets may elicit T cell responses individually and accumulatively. We therefore used the IEDB population coverage tool ^41^, which employs global HLA allele prevalence data to predict the percentage of individuals in a regional population to respond to a given epitope set. Starting with each SARS-CoV-2 peptide and predicted HLAs individually, we find considerable coverage of 55.32% for “LLLD*”, while “VQID*” exhibits the lowest predicted coverage of 7.09% (Fig 6D).

Similarly to a previous approach by Ahmedid et al ^42^, we set out to predict the accumulated global population coverage of the set. We found that 8 peptides collectively produce >90% global coverage, while the entire set is predicted to elicit T cell responses in 92.93% of the global population (Fig 6E). Regionally, Europe and North America exhibited the highest predicted coverage (Fig 6F). Of note, Africa and Asia also exhibited high predicted coverage. Central America (defined as Guatemala and Costa Rica) exhibited low coverage of 7%. It is unclear why, and further investigation is necessary to produce a peptide set with high coverage in these countries.

Overall, we identified a set of 10 SARS-CoV-2 immunogenic peptides, each highly conserved across coronavirus strains, which collectively provide global population coverage of ~93%. We believe that this is an encouraging insight in the search for pan-coronavirus T cell targets, and additionally propose these as top candidates for cross-protective immunity.

## Discussion

Our work demonstrates that T cells specific to SARS-CoV-2 peptides with high similarity to HCoV pMHC can be expanded from naïve individuals, and that these cognate public TCRs are also observed in a subset of recovered COVID-19 patients. This finding firstly suggests that SARS-CoV-2-unexposed individuals could mount T cell responses to HCoVs that - due to peptide similarity - could be cross-reactive with SARS-CoV-2 antigens. Furthermore, we propose that while COVID-19 disease appears to primarily direct responses against SARS-CoV-2-private peptides, patients with certain HLA alleles (e.g HLA-B*07:02, -C*07:02, - A*03:01) may be more likely to possess sCoV-2-HCoV cross-reactive CD8+ T cells. It is therefore plausible that SARS-CoV-2 naïve individuals with certain HLAs may be at lower risk of severe disease - or experience augmented vaccine responses - if previously exposed to endemic coronaviruses, however a direct link to pre-existing immunity requires further investigation.

Indeed, our analysis indicates that after SARS-CoV-2 infection, a subset of individuals has memory T cells that primarily recognize sCoV-2-HCoV peptides. In these convalescent patients, it is unclear whether infection itself, and/or prior exposure to HCoVs are driving this subset of individuals to select for these peptides. There is conflicting evidence surrounding the existence of memory SARS-CoV-2 cross-reactive CD8+ T cells in unexposed individuals^15,16,40^, and a limitation of our work is that we could not to provide a direct link to pre-existing immunity, because from healthy donors the MIRA dataset only evaluated expanded naïve T cells and did not examine anti-viral efficacy of the responding T cells. Indeed, although we cannot determine the cause or timeframe of this selection of sCoV-2-HCoV peptides in this subset of individuals, the potential implications are interesting. It is plausible that these patients may exhibit more robust protection against SARS-CoV-2 variants, HCoVs or even future emerging coronavirus strains. Future work should explore any immunity benefit of infection-induced cross-reactive T cell responses, and in addition, it will be interesting to examine whether vaccination against SARS-CoV-2 can induce T cell memory that is cross-reactive with SARS-CoV-2 variants and/or wider coronaviruses in such individuals. Furthermore, by our identification of a set of 10 potentially cross-reactive peptides with broad population coverage, it is possible that these peptides could be employed to test which patients exhibit cross-reactive phenotypes e.g., after vaccination with relevant antigens.

More broadly, data are beginning to demonstrate distinct vaccine-induced responses linked to differential patient exposure to SARS-CoV-2 ^3,4^. In turn, it is possible that COVID-19 vaccine boosted cross-reactive immune responses may influence vaccine-induced protection ^7^. Indeed, it will be important to explore whether COVID-19 vaccination can boost any infection-induced cross-reactive T cell memory, and whether this affects robustness of protection from SARS-CoV-2 variants or wider coronaviruses.

SARS-CoV-2 reactive CD8+ T cells have been associated with milder disease ^43^, and as previously mentioned, conflicting evidence has recently emerged regarding the presence of pre-existing CD8+ T cells in unexposed patients. Nguyen et al^16^., found that SARS-CoV-2 CD8+ T cells in Australian pre-pandemic samples, including those recognising the immunodominant HLA-B*07:02-SPR* complex, predominately displayed a naïve phenotype, indicating a lack of pre-existing memory conferred by HCoV. In contrast, Francis et al^15^., found that ~80% of unexposed individuals carrying HLA-B*07:02 show a pre-existing CD8+ T cell response to HLA-B*07:02-SPR*. Francis et al argue that these pre-existing memory pools are likely induced by prior exposure to HCoV, and that only a subpopulation of individuals carrying specific HLA would possess such memory T cells. Our work is consistent with a subset of COVID patients enriched for carrying HLA-B*07:02, and observed that in these patients, their public T cells respond primarily to shared peptides. Despite not providing a link to memory vs naïve responses, we build upon existing work by proposing additional alleles which may be carried by individuals who possess cross-reactive T cells, as well as those which appear depleted or absent in these individuals. Few studies have examined associations between HLA type and COVID disease or its severity ^15,44,45^. Nevertheless, the emerging picture is indicating that HCoV-SARS-CoV-2 cross-reactivity is conditioned by HLA genotype. Together, we provide a landscape of TCR-pMHC interactions (all TCR-pMHC interactions used in the analyses are found in Supplementary Data File 8) which may be involved in HCoV-SARS-CoV-2 cross-reactivity and provide a framework for further anti-viral mechanistic studies.

Although our study provides a map of shared and private SARS-CoV-2 peptides to date and offers the extent to which one may expect CD8+ T cells cross-reactivity between HCoVs and SARS-CoV-2, a limitation is that for cross-reactivity insights, we had to limit ourselves only on CD8+ T cells for which both peptides and their cognate TCRs information were available. Additionally, our approach for identifying homologous sequences seems to work better for MHC class I peptides that are considerably shorter in length than their class II counterparts. With a more suitable metric for longer peptides, one may substantiate our insights for class II.

Our metric for discriminating shared and private peptides is based on three factors: 1) sequence homology at 50%, 2) physicochemical similarity of 75% and 3) that both source and target peptides must be presented by the same HLA. Of these three, 50% sequence homology may seem too relaxed. In support of our use of this threshold we note that: a) factors 2 and 3 are additionally applied to compensate for this, b) we have checked our results with 70% sequence homology and observed that main conclusions are robust, 3) as this map is suggested for further functional validation we favour minimizing false negatives at the cost of potential false positives.

Through examining the potential for cross-reactivity between SARS-CoV-2 and HCoV strains, we have predicted that a set of 10 highly conserved immunogenic peptides could mount CD8+ T cell responses in 99% of the global population. These peptides have been reported previously in *in silico* and experimental work ^46-50^ however to our knowledge their large accumulated global population coverage has not yet been reported. Some of these peptides exhibit similar population coverage although with different HLA profiles, therefore it may be possible to tailor a smaller set of peptides to specific regions of interest (based on local HLA frequency), thus maximising coverage with a minimal set of peptides. Our work firstly identifies these peptides as top candidates for cross-reactivity. Secondly, we propose that their high conservation across strains may be of interest as pan-coronavirus targets, to assist ongoing work in search of mitigation strategies to reduce the threat from mutant variants or emerging coronaviruses ^51-53^.

A complex facet of severe COVID-19 disease and its diverse clinical manifestation is immunopathogenesis. Indeed, exacerbated immune responses including cytokine storm are a primary clinical characteristic in severe COVID-19 patients. Aberrant transcriptional programming has been observed in response to SARS-CoV-2 ^54^, characterised by a failure of type-1 and −3 interferon responses and simultaneous high induction of chemoattractants. While the growing evidence for pre-existing HCoV cross-reactive memory T cell responses may simply translate into an immunity benefit in some patients, in concert with data from MERS and SARS-CoV-1, there are considerable evidence that cross-reactive T and B cell responses may on the other hand be involved in immunopathology with SARS-CoV-2.

Venkatakrishnan et al.,^55^ identified peptides that are identical between SARS-CoV-2 and the human proteome. Their work demonstrates that the genes giving rise to these peptides are expressed in tissues implicated in COVID-19 pathogenesis. Our work expands their insights, by identifying SARS-CoV-2 peptides that are experimentally confirmed to be immunogenic, with high similarity to the human proteome. Consistent with their conclusions, we find similarity of immunogenic SARS-CoV-2 peptides to human genes e.g., CCL3, CCL31 and CD163. These insights are of particular interest given the elevated cytokine and chemokine responses in severe COVID patients. More broadly, there is evidence that viral antigens that are structurally similar to self-antigens can be involved in inducing autoimmunity via molecular mimicry ^8^. For these reasons, we propose these peptides as candidates which may exhibit immunopathological or autoimmune associations.

In conclusion, we have employed an *in-silico* approach to examine the evidence surrounding cross-reactive SARS-CoV-2 CD8+ T cell responses. We observed a set of SARS-CoV-2 candidates with high similarity to the human proteome and suggest investigation into whether they provoke immunopathology. We have also provided evidence of CD8+ T cell cross-reactivity, not only to an extent which indicates that naïve individuals could mount cross-reactive responses to SARS-CoV-2 and common-cold coronaviruses, but we also found that SARS-CoV-2 infection induces CD8+ T cell responses against peptides with high similarity to HCoV in some COVID-19 patients. We build upon existing evidence that such cross-reactivity is conditioned by presence of specific HLA alleles and envision that the insights presented here are leveraged to explore whether these potentially cross-reactive T cells and cognate pMHCs influence COVID-19 disease heterogeneity, vaccine-or infection-induced protection from SARS-CoV-2 and its emerging variants of concern.

## Acknowledgments

We greatly acknowledge conversations and guidance from Dr Mikhail Shugay (ITM, Moscow), and Dr Giorgio Napolitani (KCL, London).

## Author contributions

HK conceived, designed and supervised the project. PB performed computational analyses with insights from CL and MPP and AA. HK and PB interpreted the results. AS, GO assisted design, interpretation and supervision. ROB, JW commented on manuscript. HK, AS funded the project. HK, PB, wrote the manuscript with contributions from CL, MPP and AA, AS and GO.

## Methods

### Data Processing and Analysis

All data processing and analysis was performed using the R plugin for Pycharm 2020, in either R 4.0.3 or 4.0.1.

### Curating a pool of SARS-CoV-2 class I and II peptides

Human immunogenic and non-immunogenic SARS-CoV-2 peptide data were gathered from the both the IEDB and the Virus Pathogen Resource (VIPR) (accessed 11-02-2021). ‘T cell’ assay, ‘Human’ host and SARS-CoV-2 organism options were selected. If an observation was found in both datasets, the one from the IEDB was retained. Protein names were cleaned and standardised where possible. Immunogenic peptides not observed in either the IEDB or VIPR were also gathered from the ‘MIRA’ dataset which maps cognate TCRs and SARS-CoV-2 peptides.

### Retrieval of Coronavirus Proteome Sequences

NCBI reference genomes were gathered for OC43 (https://www.ncbi.nlm.nih.gov/nuccore/1578871709/), HKU1 (https://www.ncbi.nlm.nih.gov/nuccore/NC_006577.2), 229E (https://www.ncbi.nlm.nih.gov/nuccore/NC_002645.1), NL63 (https://www.ncbi.nlm.nih.gov/nuccore/49169782/), and SARS-CoV-2-Wuhan (https://www.ncbi.nlm.nih.gov/nuccore/NC_045512.2).

### MHC Presentation Prediction

Antigen presentation by MHC class I was predicted using NetMHCpan v4.1 against HLA-A*0101, 0201, 0301, 2402, HLA-B*0702, 4001, 0801, and HLA-C*0702, 0401, 0701 alleles. Antigen presentation by MHC class II was predicted using netMHCIIpan against the most common sets of alleles found in the IEDB, for which this model can make predictions to. The alleles are: DRB1-0101, 0102, 0301, 0401, 0402, 0402, 0404, 0701, 0801, 0901, 1001, 1101, 1104, 1201, 1202, 1301,1302, 1303, 1401, 1406, 1501, 1502, 1601, 1602, DRB3_0101, 0202 and DRB5_0101, 0102. Peptides with a rank score <=2.0 were classified as binders.

### HLA ligand enrichment analysis for SARS-CoV-2 proteins

To provide reasonable statistical inference, we only examined proteins longer than 100 amino acids. To compute enrichment or depletion, we followed the approach by Karnaukhov et al. First, we predicted using netMHCpan v 4.1 the number of ligands *Ni* of length *l* from each SARS-CoV-2 protein *i* which adheres to the criteria. Probability of a HLA allele presenting a peptide was computed as the average number of ligands per allele:

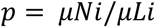

where Li is the corrected protein length (length of protein - *l*), and μ denotes the average over the assessed SARS-CoV-2 proteins. It follows that the probability of observing a given number of ligands from each SARS-CoV-2 protein is computed using the binomial distribution as:

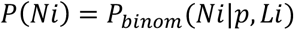

The logs odds ratio (enrichment or depletion) is calculated as:

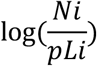

### Discriminating Shared and Private SARS-CoV-2 Peptides

To compare a SARS-CoV-2 peptide *a*, of length *N* to a proteome of interest, all possible linear peptides of length *N* were generated from said proteome. This can be thought of as scanning along the proteome of interest with a step size of 1, generating all peptides of length *N*. The deriving protein was recorded. Three metrics - which all must be satisfied - were used determine whether a peptide is considered shared with HCoV or private to SARS-CoV-2. We below describe each metric, and then explain the three thresholds which all must be achieved for a peptide to be classified as ‘shared.’.

Firstly, once all peptides from the proteome of interest of length *N* are generated, a similarity index we call the ‘MatchScore’ is calculated for each pairwise comparison. This metric is charged with assessing physicochemical similarity between two peptides of interest. For each SARS-CoV-2 peptide, the highest ‘MatchScore’ against each HCoV protein is retained and the rest are discarded. To calculate the ‘MatchScore’, we employ the method designed by Bresciani et al ^24^. Briefly, for two peptides *a* or *b* of length N, the similarity score is given as;

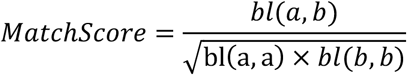

where bl(a,b) is the BLOSUM62 score for peptide *a* vs *b*, and bl(a,a) is the BLOSUM62 score for peptide *a* vs *a*, etc. BLOSUM62 local-global alignment scores (local or global would produce the same score for a pairwise alignment of lengths N vs N) were computed using the pairwiseAlignment function from the R package Biostrings, with high gap penalties (opening and extension of both 100). The MatchScore function produces a score where 1 reflects an exact match, i.e no mismatches in two sequences, and 0 reflects high dissimilarity.

#### Criteria 1: A shared peptide and its HCoV match must have a MatchScore of >0.75

The second metric is based on sequence homology between two sequences, essentially reflecting the proportion of amino acid positions in the SARS-CoV-2 peptide, which are conserved in the HCoV match. This is calculated as:

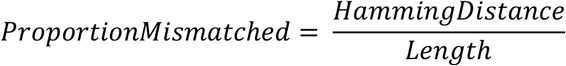

where ‘*HammingDistance*’ is the hamming distance between two peptides of interest, which calculates the number of different positions, and ‘*Length*’ is the length of the compared peptides.

#### Criteria 2: The *ProportionMismatched* between a shared peptide and its HCoV match must be < 0.5 (50%)

Naturally, the inverse of this is true, in that at least 50% amino acid conservation between a SARS-CoV-2 peptide and a HCoV match must be observed for the peptide to be considered ‘shared’.

The third metric is based on predicted presentation by HLA of the SARS-CoV-2 peptide and its HCoV match.

#### Criteria 3: Both the SARS-CoV-2 peptide and its HCoV match must be predicted to bind at least one common HLA allele

All three criteria must be satisfied for a SARS-CoV-2 peptide to be classified as a shared peptide and also for a match from HCoV to be considered a homologous match. *doParallel* and *foreach* functions were used to parallelise the processing.

### Sequence Similarity with the Human Proteome and Human Microbiomes

Here, the same similarity criteria were employed as in the previous HCoV section. However, in contrast with HCoV comparison, due to the size of the human proteomes and microbiomes, the best match against the whole proteome is retained. *doParallel* and *foreach* functions were used to parallelise the processing.

### Gathering human proteome sequence

The human proteome was downloaded in fasta format from UniProt https://www.uniprot.org/proteomes/UP000005640

### Gathering human microbiome sequences

Human gut and airways microbiomes were downloaded from the HMP Data Analysis and Coordination Center http://www.hmpdacc.org/HMRGD. The complete set of genomes were downloaded in fasta format in ‘Protein multifasta (PEP) format’. For gut, the body site was specified as ‘gastrointestinal tract’. 457 and 50 gut and airway microbiota were available respectively.

### Human gene sets with sequence similarity to SARS-CoV-2 immunogenic peptides

The SARS-CoV-2 peptides of lengths 9 and 10 with a similarity score to the human proteome in the top 10 percentile were gathered. Only predicted binders (see MHC presentation prediction) were retained.

### CD8+ T cell cross-reactivity maps using IEDB receptor data

The entire IEDB receptor data for SARS-CoV-2 peptides was downloaded. Bipartite graphs were generated using *iGraph* and *Matrix* libraries in R. Bipartite graphs were projected into one-mode graphs using the *bipartite_projection* function. All graphs were exported from *iGraph* into Cytoscape v3.82 using the R function *createNetworkFromIgraph* from package *RCy3*. From Cytoscape, ‘.graphml’ files were exported and opened with Gephi. Gephi was used to finalise the diagrams and improve visual aesthetics. Either ‘ForceAtlas’ or ‘Fructerman-Reingold’ templates were used. Gravity and repulsion parameters were altered to improve visual aesthetics.

### CD8+ T cell CDR3 Kmer Enrichment

R Package *immunarch* ^56^was used to compute Kmer (K=5 in this case) statistics for CDR3 sequences, and to visualise enrichment. See https://immunarch.com/articles/web_only/v9_kmers.html for full details.

### Gathering clinical and TCR repertoire data for COVID-19 patients and healthy subjects

The COVID-19 MIRA dataset (>160k high-confidence SARS-CoV-2-specific TCRs) was downloaded from https://clients.adaptivebiotech.com/pub/covid-2020 with corresponding sample metadata. These data contain TCR repertoire data mapped to SARS-CoV-2 epitopes from 5 patient cohorts, including COVID convalescent patients and healthy subjects with no known exposure to SARS-CoV-2. Only convalescent patients and healthy subjects were used in the analysis due to low numbers of subjects for other cohorts.

### Networks of TCRs recognising Shared and/or Private Peptides

A public TCR is defined as a CDR3 sequence and V and J gene which is observed in more than one patient in the MIRA dataset. All graphs were first generated using *iGraph* in R, exported to Cytoscape using the *createNetworkFromIgraph* function in the *RCy3* package. From cytoscape, all graphs were exported as .graphml files and read into Gephi. In Gephi, either ‘ForceAtlas’ and ‘Fruchterman-Reingold’ templates were used. In almost all cases, gravity and repulsion parameters were adjusted to improve visual aesthetics.

### Estimating population coverage of SARS-CoV-2 peptides with high conservation to three or more HCoV

We followed the approach by Ahmedid et al^42^. Population coverage is an estimate of the proportion of individuals in a given population that may mount a T cell response against a peptide. Population coverage is predicted based on HLA alleles for each immunogenic peptide as predicted by netMHCpan 4.1, leading to individual population coverage of a peptide. To predict accumulated coverage, we began with the peptide with the highest individual coverage “FVDG*”, and incrementally added a peptide and predicted accumulated coverage. The population coverage of a set of peptides (i.e accumulated coverage), is defined as the proportion of individuals able to mount a T cell response to at least one peptide in the set. Python code for the IEDB tool to compute the population coverage was downloaded from http://tools.iedb.org/population/download on 24-11-20.

## Supplementary

**Supplementary Figure 1:**
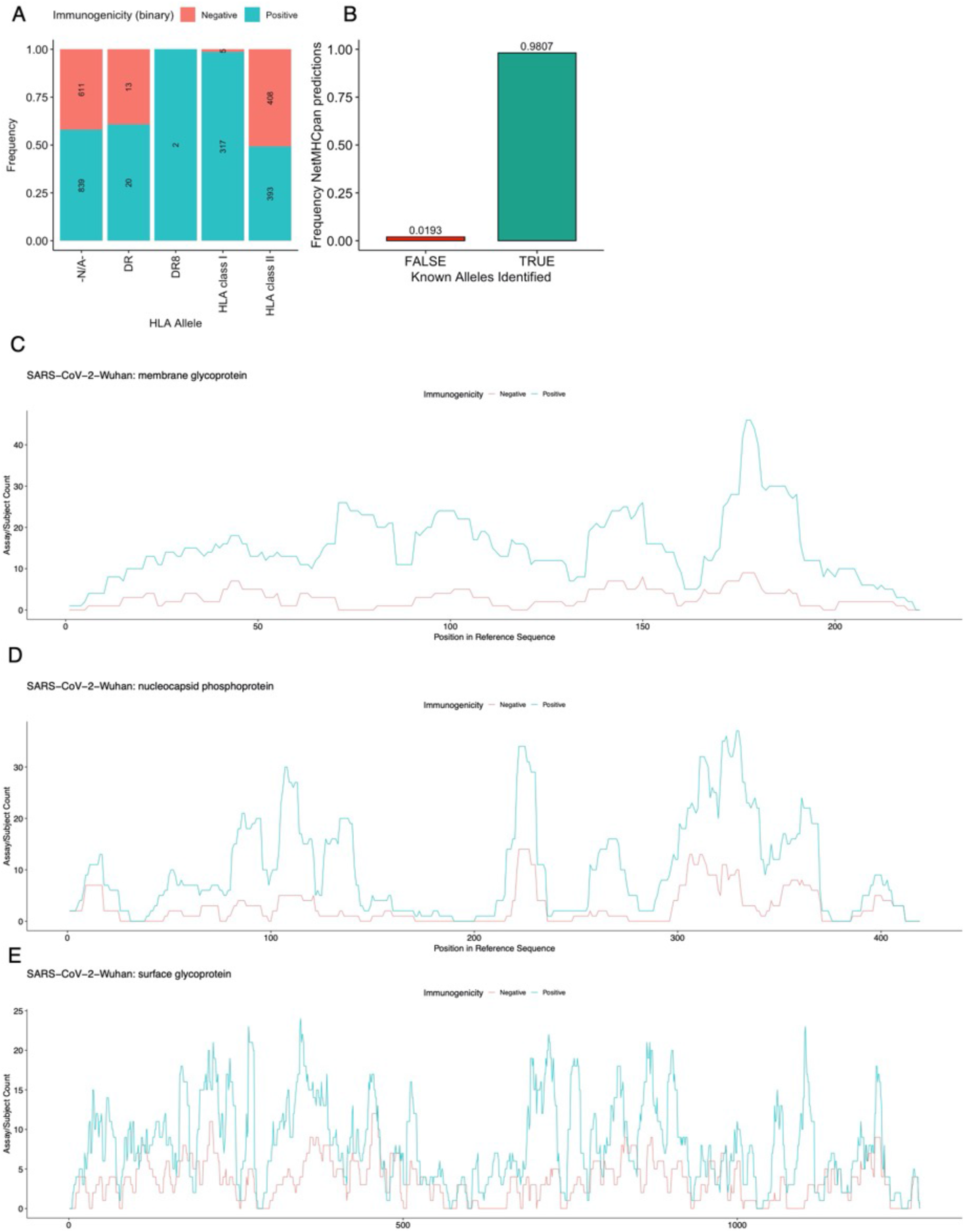
A-B) Barplots showing A) the frequency of immunogenic epitopes amongst HLA labels that do not designate a specific allele. B) the frequency of antigen presentation predictions where the correct allele for the SARS-CoV-2 peptide (where known) was identified. C-E) Line plots showing the immunogenic regions of the most immunodominant SARS-CoV-2 proteins, C) membrane glycoprotein, D) nucleocapsid phosphoprotein, E) ‘spike’ surface glycoprotein.

**Supplementary Figure 2:**
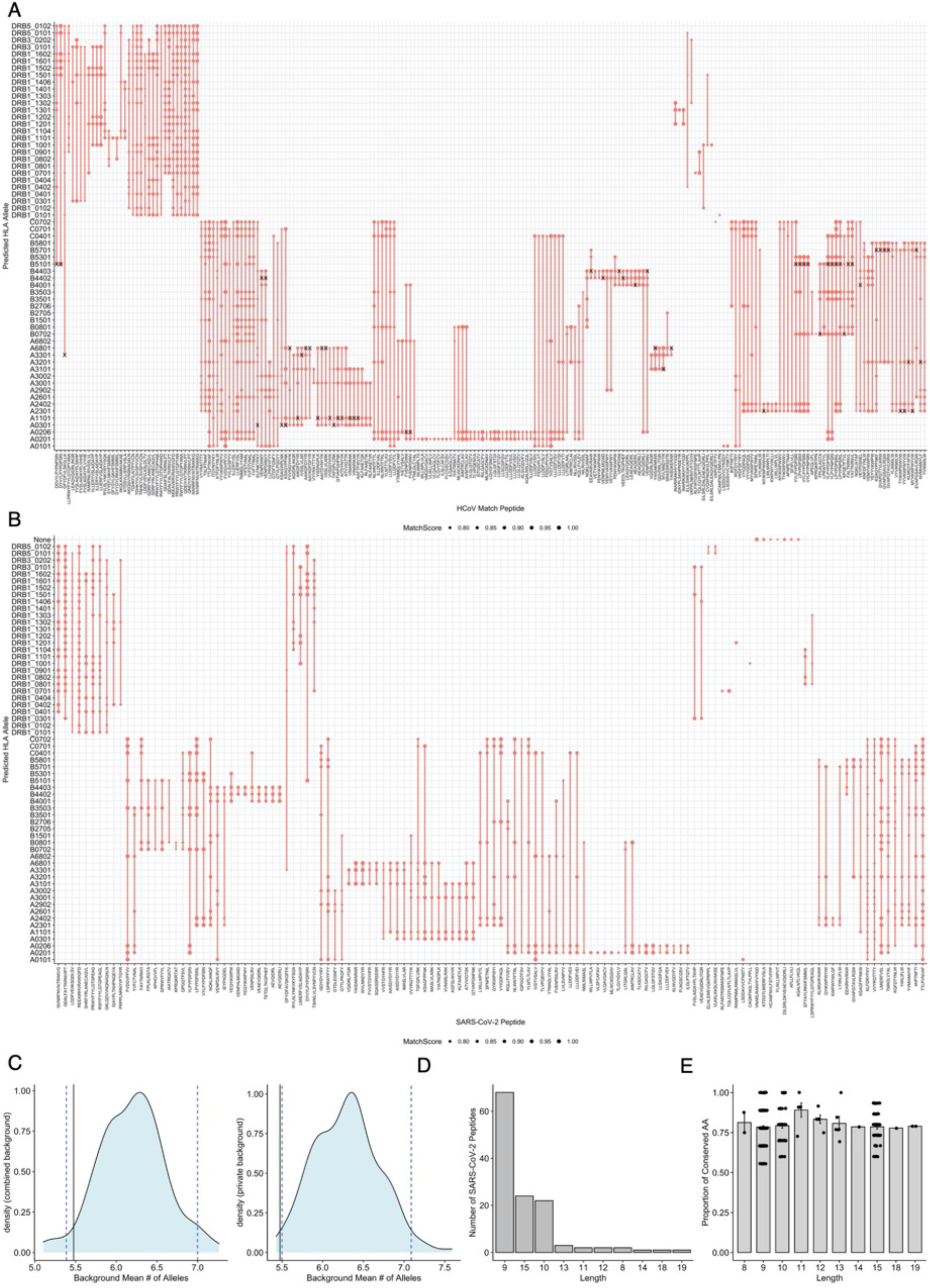
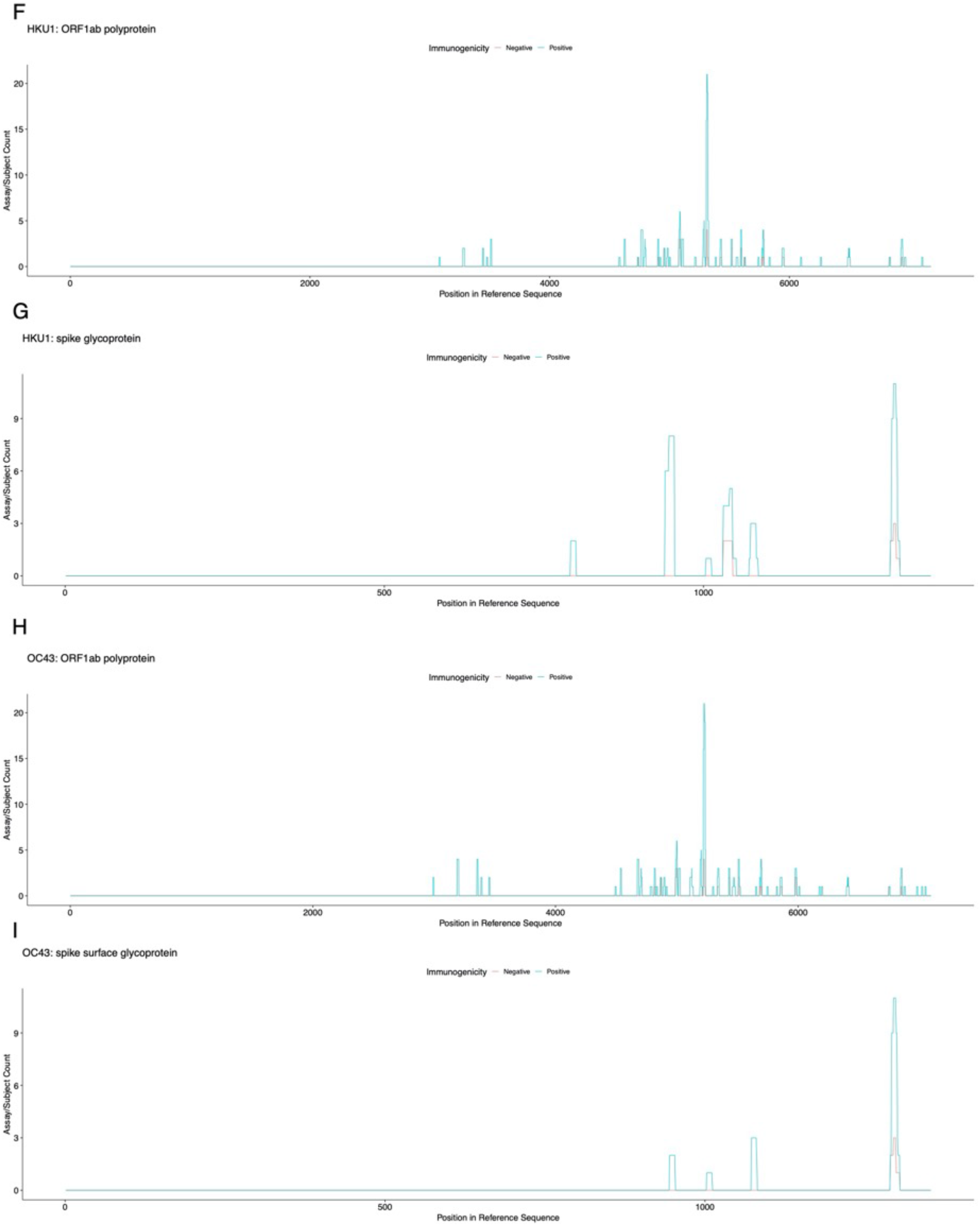
**A)** A dot plot showing predicted HLA alleles for peptides from HCoV with high-similarity to immunogenic SARS-CoV-2 peptides. Size of the point reflects the similarity score to the corresponding SARS-CoV-2 peptide. Lines link predicted alleles. An ‘X’ indicates – where possible - if there is experimental evidence that the corresponding SARS-CoV-2 peptide binds the predicted HLA allele for the HCoV match. B) a dot plot showing predicted HLA alleles for the 129 shared SARS-CoV-2 immunogenic peptides. C) Density plots showing the mean number of alleles for which shared peptides are predicted to bind (solid black line), compared with a background distribution generated by randomly shuffling the dataset using both private and shared peptides (left) or private peptides only (right). Dashed lines show +/- 2 standard deviations from the mean. D) Bar plot showing the distribution of lengths for 129 shared SARS-CoV-2 immunogenic peptides. E) Bar plot showing the distribution of amino acid conservation between the 285 matches and the 126 SARS-CoV-2 shared peptides. F-I Line plots showing the HCoV protein regions which produce immunogenic HCoV-CoV-2 shared peptides from F) HKU1 ORF1ab polyprotein, G) HKU1 spike glycoprotein, H) OC43 ORF1ab polyprotein, I) OC43 spike glycoprotein.

**Supplementary Figure 3:**
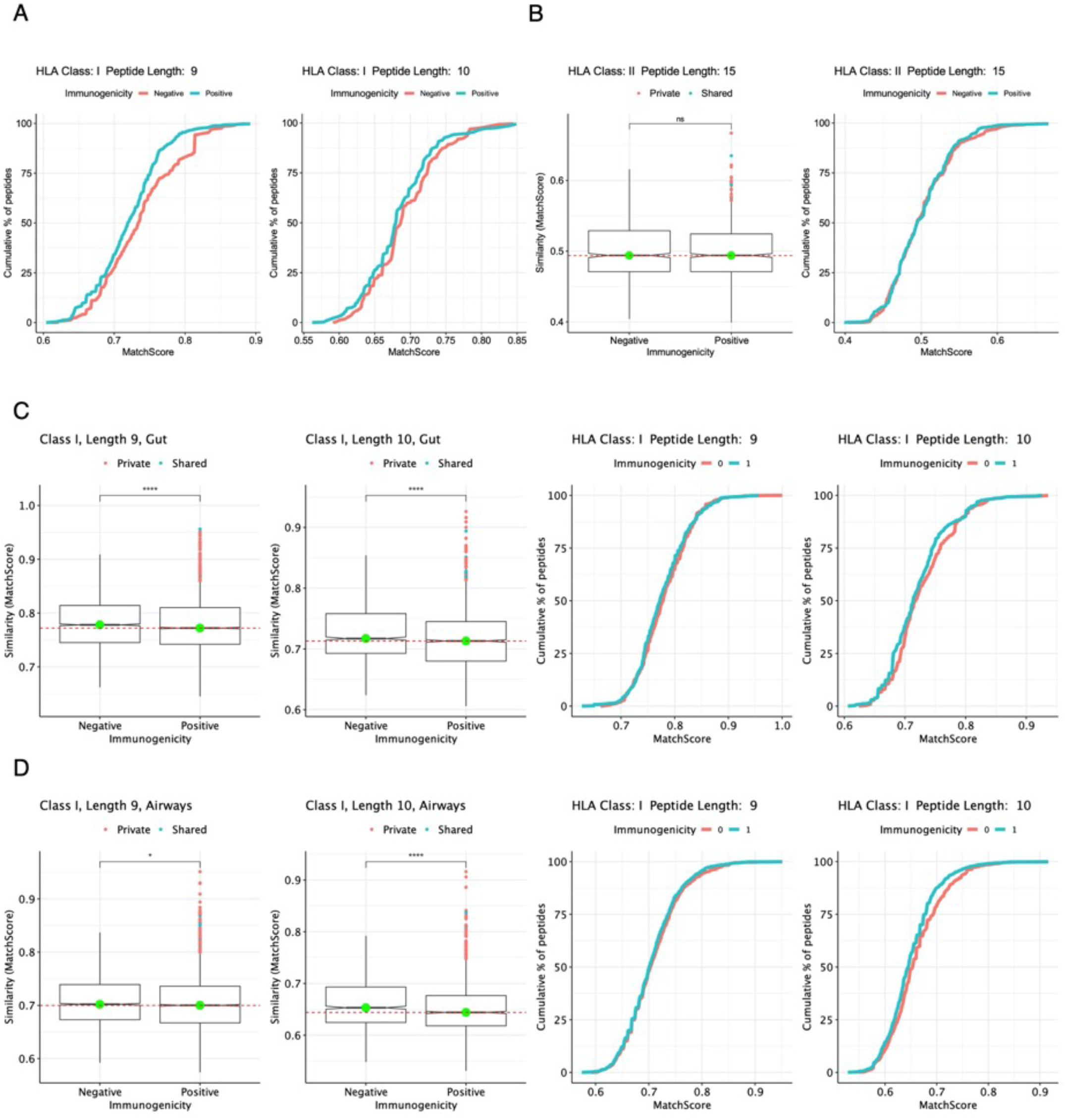
A) Empirical cumulative distribution plots for HLA class I peptides of length 9 and 10, showing the cumulative % of peptides to reach a specific MatchScore, color labelled by immunogenicity status. B) A notched boxplot and empirical cumulative distribution plot showing the similarity -as evaluated by the MatchScore - of nonimmunogenic and immunogenic class II SARS-CoV-2 peptides with sequences derived from the human proteome of length 15 (immunogenic n=955, nonimmunogenic n=953 peptides). C-D) Notched boxplots and empirical cumulative distribution plots showing the similarity of nonimmunogenic and immunogenic SARS-CoV-2 peptides with sequences derived from gut C) and airway microbiomes D).

**Supplementary Figure 4:**
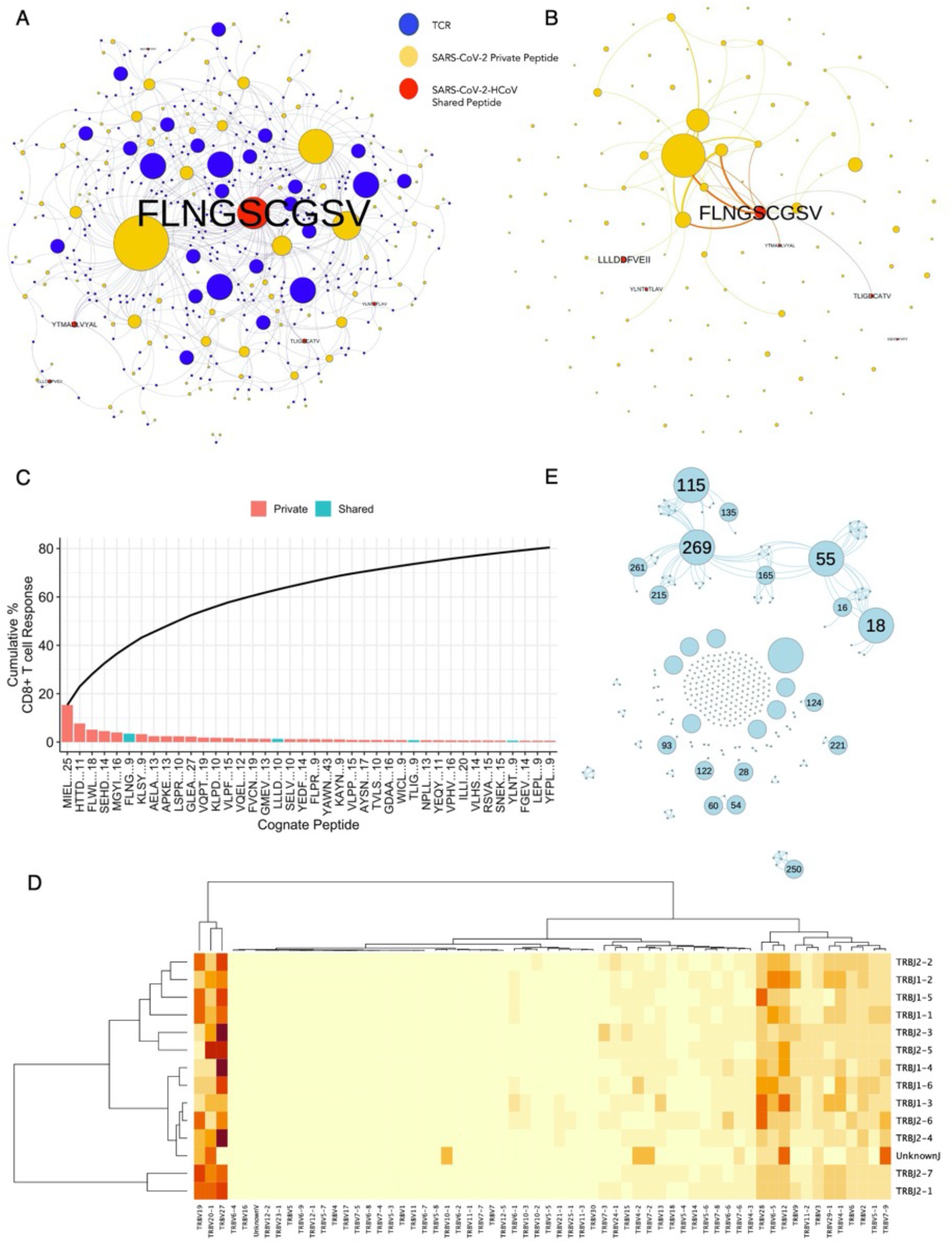
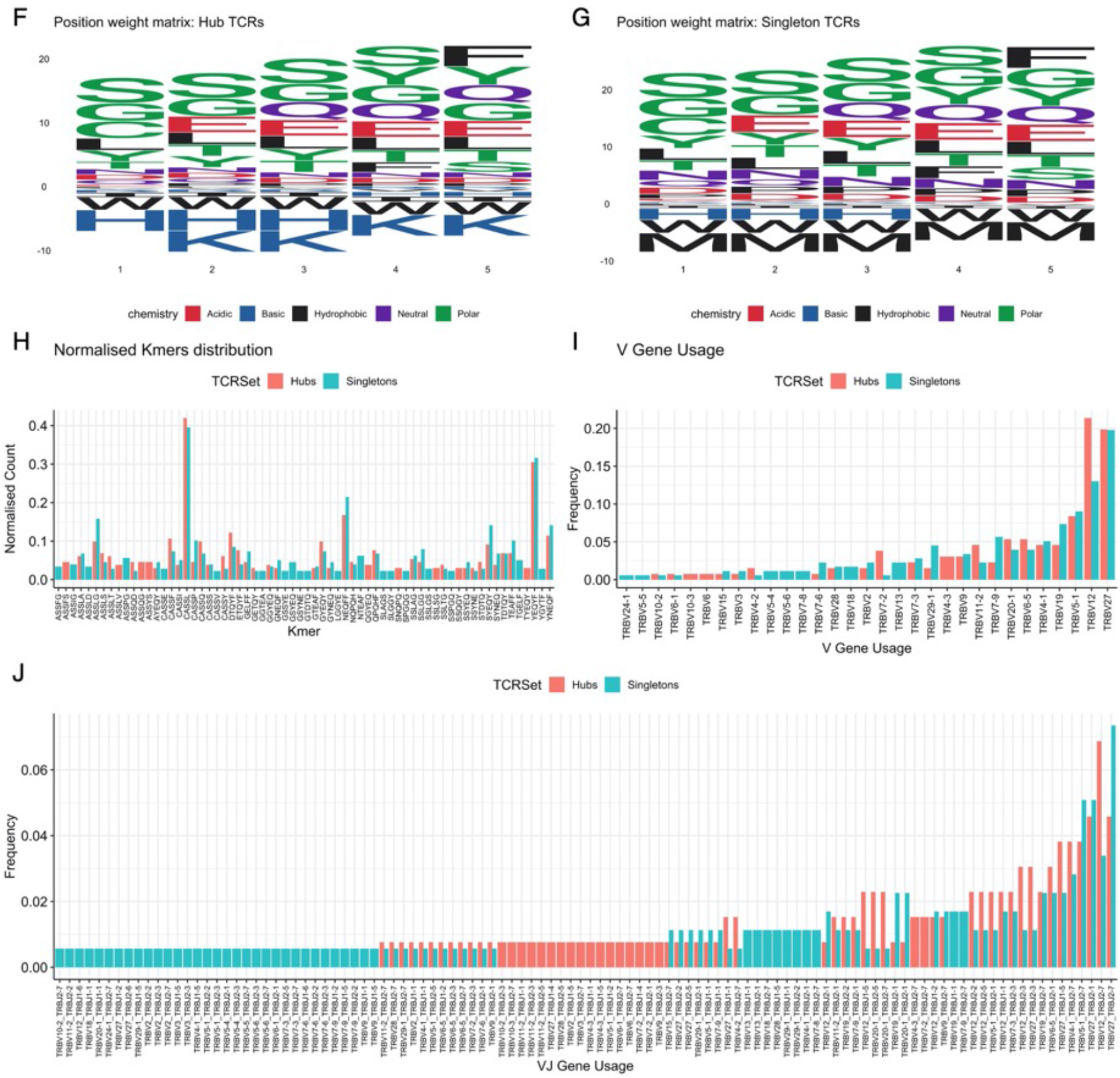
CD8^+^T cell cross-reactivity against SARS-CoV-2. A) A bipartite network graph depicting the interactions with SARS-CoV-2 immunogenic peptides (SARS-CoV-2-shared are colored red, -private are coloured yellow) and their cognate TCRs (blue). Node size represents the degree of connectivity. B) A one-mode network graph showing shared and private SARS-CoV-2 peptides. An edge between a peptide node demonstrates that a peptide is recognized by the same TCR. Node size reflects the degree of connectivity. C) A barplot showing the top 80% of the cumulative SARS-CoV-2-specific CD8+ T cell response and the peptides recognized, as per the cognate TCR dataset from the IEDB. D) A heatmap showing common V and J gene usage for SARS-CoV-2 specific TCRβ sequences. E) A one-mode network graph showing the common specificity of SARS-CoV-2 specific TCRs. Each node is a TCR, and an edge reflects whether two TCRs recognize the same peptide. Node size reflects the number of peptides recognized by a TCR. F) A sequence logo plot visualizing the position weight matrix for CDR3b 5mers in the IEDB dataset, for “Hub TCRs”, those with considerable common-specificity. G) A sequence logo plot visualizing the position weight matrix for CDR3b 5mers in the IEDB dataset, for “Singletons”, those recognizing only one unique SARS-CoV-2 peptide H) A barplot contrasting the Kmer distribution for “Hub” and “Singleton” TCRs. The count is normalized by the number of TCRs in each group (Hub or Singletons). I) A barplot contrasting the V gene usage of “Hub” and “Singleton” TCRs. Y axis shows the frequency to which the V gene is used in each group. J) A barplot contrasting the V-J gene usage of “Hub” and “Singleton” TCRs. Y axis shows the frequency to which the V-J gene combination is used in each group.

**Supplementary Figure 5:**
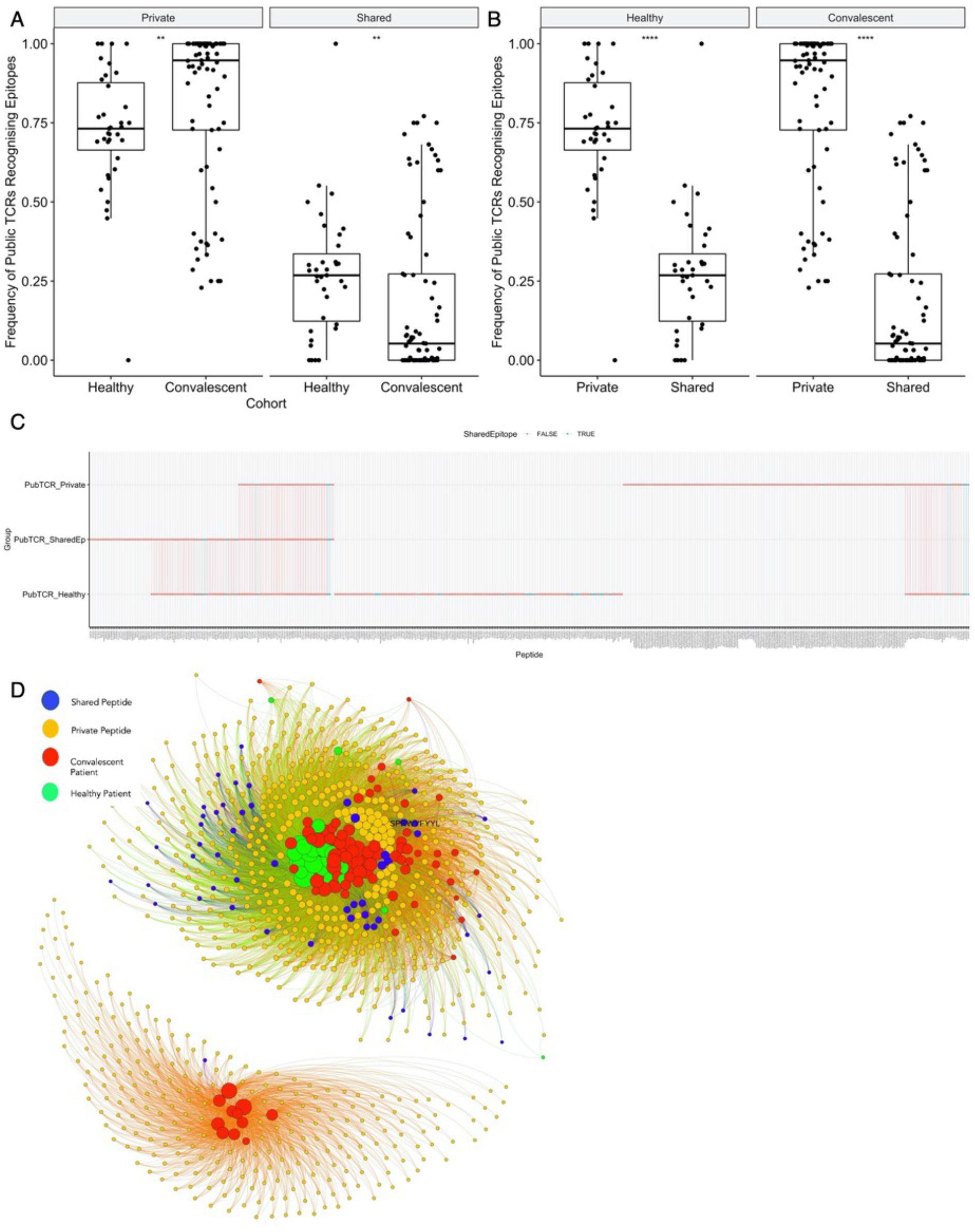
A) Boxplot showing for each patient, the frequency of public TCRs recognizing SARS-CoV-2 private or SARS-CoV-2-HCoV shared peptides, grouped by Private/Shared epitopes and contrasting patient cohort. B) Boxplot showing for each patient, the frequency of public TCRs recognizing SARS-CoV-2 private or SARS-CoV-2-HCoV shared peptides, grouped by patient cohort and contrasting Private/Shared epitopes. C) Line graph showing the peptides recognized by public TCRs in different patient cohorts: Healthy patients (PubTCR_Healthy) (n=12), COVID convalescent patients whose public TCRs predominately recognize Private (PubTCR_Private) or Shared (PubTCR_SharedEp) peptides (both n=12). D) A bipartite graph showing the common-specificity of private TCRs recognizing SARS-CoV-2 shared and private peptides. An edge between a patient and a peptide is observed if a patient posseses a private TCR recognizing that peptide.

**Supplementary Figure 6:**
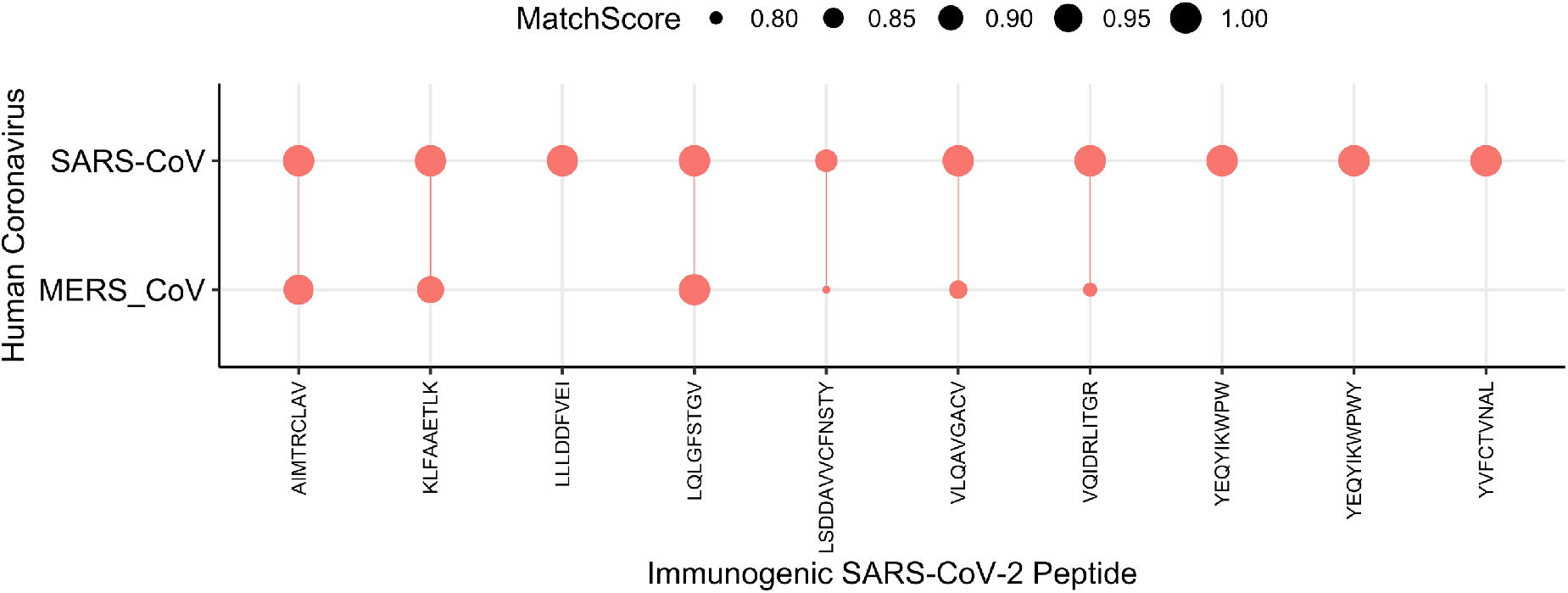
**A)** A dot and line plot showing each SARS-CoV-2 peptide on the x-axis. A dot shows a high similarity match to MERS or SARS-CoV. The size of each point reflects the MatchScore, i.e the similarity metric.

**Table S1:**
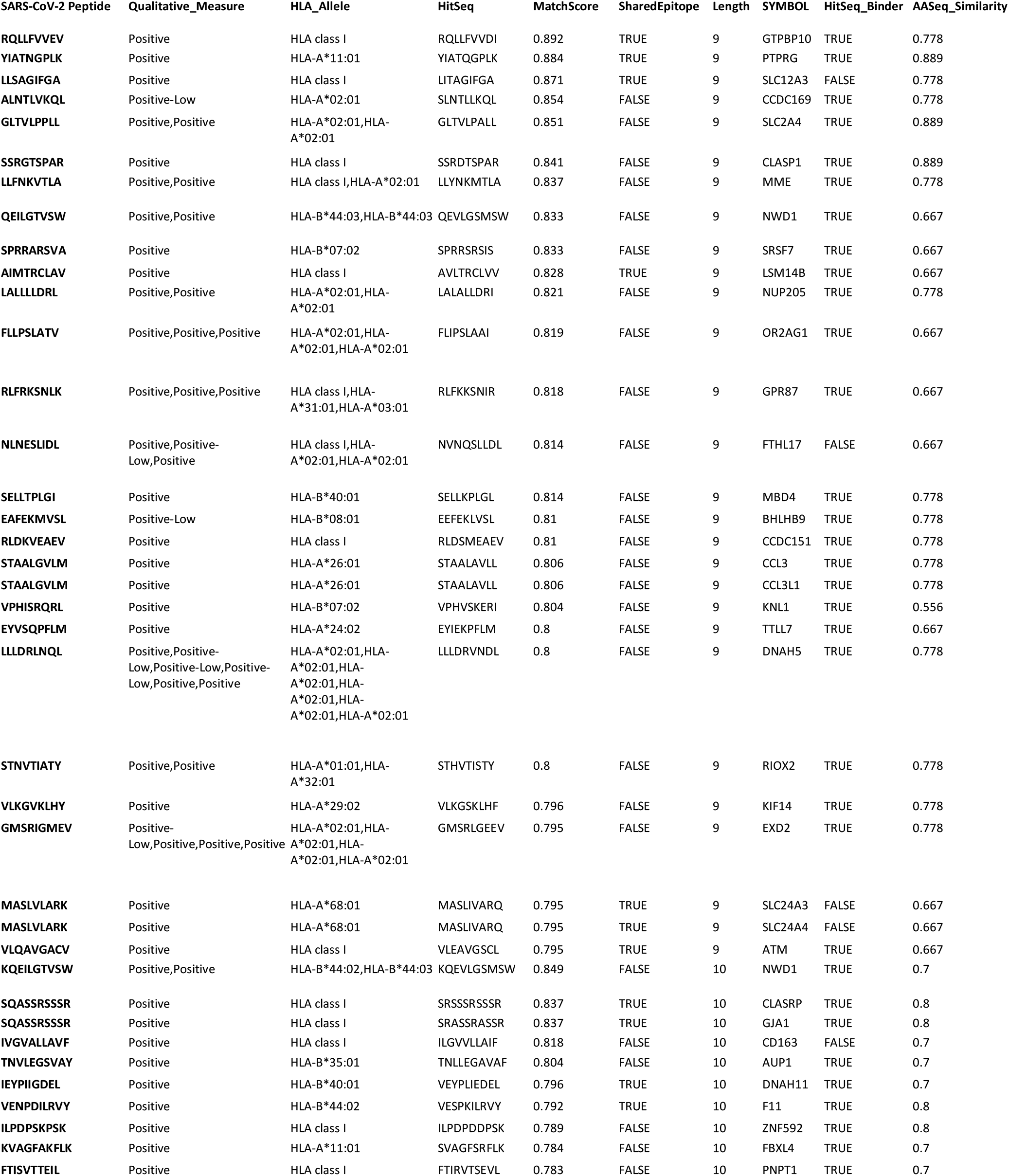
Peptides identified with high similarity to human proteome. HitSeq shows the match from the self proteome. SharedEpitope shows whether the peptide is a sCoV-2-HCoV shared peptide or not. SYMBOL shows the gene from which the hit peptide is derived. HitSeq_Binder shows whether the hit peptide is predicted to bind an HLA allele. AASeq_Similarity shows the proportion of amino acids conserved between the SARS-CoV-2 peptide and the human proteome match.

**Table S2:**
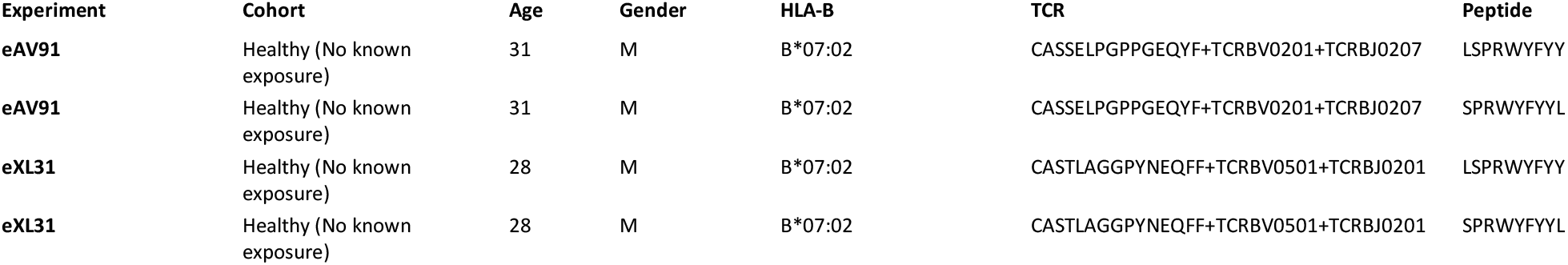
Two previously reported private TCRs are identified in additional HLA-B*07:02+ individuals at beta chain resolution, indicating these are cross-reactive public TCRs.

## Supplementary Data File description

1: Data file containing each of the 126 SARS-CoV-2 peptides which map to 285 targets from HCoV.

2: Data file containing the full set of SARS-CoV-2 peptides which have high similarity to the human proteome.

3: Data file containing a summary of the peptides recognized By public TCRs in the PubTCR-SharedEp and PubTCR-Private groups, supplemented with those recognized in a sampled set of healthy patients. File contains a 1 or a 0 demonstrating whether a public TCR in each group of patients recognizes the peptide.

4: Data file containing the information from data file 3, but also includes information pertaining to whether key class I HLA alleles are observed in a patient with a public TCR recognizing each peptide.

5: Data file containing cohort information regarding the peptides most commonly recognized by private TCRs in the MIRA dataset.

6: Data file reporting the private TCRs which recognize the sCoV-2-HCoV shared peptides shown in data file 5.

7: Data file containing information from Figure 6, exhibiting the peptides with hits to >=3 HCoV strains. File provides detailed information regarding SARS-CoV-2 peptide and the corresponding hit to HCoV.

8: Data file contains every TCR from the IEDB and the MIRA datasets which we used in this analysis, mapped to the recognized SARS-CoV-2 peptide.

